# Divisive suppression explains high-precision firing and contrast adaptation in retinal ganglion cells

**DOI:** 10.1101/064592

**Authors:** Yuwei Cui, Yanbin V. Wang, Silvia J. H. Park, Jonathan B. Demb, Daniel A. Butts

**Author notes:** these authors contributed equally to the work (Jonathan B. Demb, Daniel A. Butts).

## Abstract

Visual processing depends on specific computations implemented by complex neural circuits. Here, we present a circuit-inspired model of retinal ganglion cell computation, targeted to explain their temporal dynamics and adaptation to contrast. To localize the sources of such processing, we used recordings at the levels of synaptic input and spiking output in the *in vitro* mouse retina. We found that an ON-Alpha ganglion cell’s excitatory synaptic inputs were described by a divisive interaction between excitation and delayed suppression, which explained nonlinear processing already present in ganglion cell inputs. Ganglion cell output was further shaped by spike generation mechanisms. The full model accurately predicted spike responses with unprecedented millisecond precision, and accurately described contrast adaption of the spike train. These results demonstrate how circuit and cell-intrinsic mechanisms interact for ganglion cell function and, more generally, illustrate the power of circuit-inspired modeling of sensory processing.

## INTRODUCTION

Neural computations in the retina are generated by complex circuits that drive the responses of ~30 distinct ganglion cell types (Baden et al., 2016; Demb and Singer, 2015; Sanes and Masland, 2015). Despite the complexity of retinal circuitry, the broad features of spike firing, in many ganglion cell types, can be described with a straightforward Linear-Nonlinear (LN) cascade model (Shapley, 2009). In this model, a linear receptive field filters the stimulus, and a nonlinear function shapes the output by implementing the spike threshold and response saturation (Baccus and Meister, 2002; Chichilnisky, 2001; Kim and Rieke, 2001). However, many aspects of ganglion cell firing deviate from LN model predictions. For example, the LN model does not capture the effect of contrast adaptation, which includes reduced gain (i.e., filter amplitude) at high contrast (Kim and Rieke, 2001; Meister and Berry, 1999; Shapley and Victor, 1978). Furthermore, the LN model does not predict firing at high temporal resolution (Berry and Meister, 1998; Butts et al., 2016; Butts et al., 2007; Keat et al., 2001; Passaglia and Troy, 2004; Uzzell and Chichilnisky, 2004), and yet precise firing represents an essential element of downstream visual processing (Bruno and Sakmann, 2006; Havenith et al., 2011; Kelly et al., 2014; Wang et al., 2010a).

To improve on the LN model, several nonlinear approaches have been proposed. The first approach describes the nonlinear function between stimulus and response as a mathematical expansion, extending from the linear receptive field (Chichilnisky, 2001) to second-order quadratic terms, using either spike-triggered covariance (Fairhall et al., 2006; Liu and Gollisch, 2015; Samengo and Gollisch, 2013; Vaingankar et al., 2012) or maximally informative dimension analyses (Sharpee et al., 2004). Such expansion terms better predict the spike train, but they are difficult to interpret functionally and with respect to the underlying circuitry (Butts et al., 2011; McFarland et al., 2013). The second approach targets specific aspects of the response, such as spike-refractoriness (Berry and Meister, 1998; Keat et al., 2001; Paninski, 2004; Pillow et al., 2005), gain changes associated with contrast adaptation (Bonin et al., 2005; Mante et al., 2008; Meister and Berry, 1999; Shapley and Victor, 1978), the interplay of excitation and inhibition (Butts et al., 2016; Butts et al., 2011), and rectification of synaptic release, associated with nonlinear spatial processing (Freeman et al., 2015; Gollisch, 2013; Schwartz and Rieke, 2011). However, each of these models primarily focuses on one type of nonlinear computation and does not generalize to explain a range of response properties.

Here we derive a novel nonlinear modeling framework inspired by retinal circuitry. The model is constrained by recordings at two stages of processing: excitatory synaptic input and spike output, recorded in mouse ON-Alpha ganglion cells. We devise a tractable model of excitatory currents that incorporates a nonlinear structure based on realistic circuit elements. In particular, we allowed for divisive suppression acting on a ganglion cell’s excitatory inputs to capture the computations implemented by presynaptic inhibition (Eggers and Lukasiewicz, 2011) and synaptic depression (Jarsky et al., 2011; Ozuysal and Baccus, 2012) at bipolar cell terminals. Ganglion cell firing, further shaped by spike generation mechanisms, could be predicted to millisecond precision. Our study establishes a unified model of nonlinear processing within ganglion cells that accurately captures both the generation of precise firing events and contrast adaptation. Similar circuit-inspired modeling could be applied widely in other sensory systems.

## RESULTS

We recorded spikes from ON-Alpha ganglion cells in the *in vitro* mouse retina while presenting a temporally modulated (<30 Hz), 1-mm spot centered on the neuron’s receptive field (Fig. 1A, *top)*. Every 10 seconds the contrast level switched between high and low. In high contrast, the stimulus evoked spike responses that were precisely timed from trial to trial (Fig. 1A, *left)*, consistent with previous work performed both *in vitro* and *in vivo* (Berry and Meister, 1998; Butts et al., 2016; Butts et al., 2007; Passaglia and Troy, 2004; Reinagel and Reid, 2000; Uzzell and Chichilnisky, 2004).

We first used a linear-nonlinear (LN) cascade model (Fig. 1B) (Chichilnisky, 2001; Hunter and Korenberg, 1986) to predict the observed responses. The “L” (linear) step of the cascade processes the stimulus with a linear “receptive field” **k**, whose output reflects the degree that the stimulus *s*(*t*) matches **k**. The “N” (nonlinear) step acts on output of the receptive field, **k** • *s*(*t*), which is scaled by a nonlinear function that could include the effects of spike threshold and response saturation. Both the linear receptive field and the nonlinearity are fit to the data in order to best predict the firing rate. The resulting receptive field had a biphasic shape at both contrasts, representing the sensitivity of the neuron to dark-to-light transitions (Fig. 1B). Furthermore, the filter had smaller amplitude at high contrast, a signature of contrast adaptation (Baccus and Meister, 2002; Kim and Rieke, 2001; Zaghloul et al., 2005).

Despite capturing the coarse temporal features of the response, the LN model could not capture fine temporal features at high contrast (Fig. 1A) (Berry and Meister, 1998; Liu et al., 2001). To precisely compare time scales of the observed data with model predictions, we performed “event analysis”, which divides the spike train into firing events separated by silence (Butts et al., 2010; Kumbhani et al., 2007). Based on this analysis, the LN model failed to predict either the SD of the first-spike in each event or the overall event duration in high contrast, but was largely successful in low contrast (Fig. 1C).

To improve on the LN model prediction, we included a refractory period (RP) following each spike (Paninski, 2004), which has previously been suggested as a mechanism for precise firing in ganglion cells (Berry and Meister, 1998; Keat et al., 2001) (see Methods). However, while the resulting LN+RP model could predict the temporal properties of the spike train at low contrast, it failed at high contrast (Fig. 1C). Thus, spike-refractoriness alone could not explain the precision at high contrast, and consequently could not predict how the response changes from low to high contrast.

### Nonlinear processing distributed across two stages of retinal processing

Because some degree of contrast adaptation is already present in a ganglion cell’s excitatory synaptic inputs (Beaudoin et al., 2007; Beaudoin et al., 2008; Kim and Rieke, 2001), we hypothesized that we might uncover the source of the nonlinear processing by directly modeling the synaptic input currents. We therefore made whole-cell patch clamp recordings on the same neurons we recorded spike responses from, and performed a similar LN analysis on excitatory synaptic currents (Fig. 1D). The LN model of the currents (Fig. 1E) – like that of the spike response – accurately predicted the observed response at low contrast, but performed relatively poorly at high contrast (Fig. 1D). To compare the precision of the LN model with the observed data, we measured the coherence between the trial-averaged response and the responses on individual trials (see Methods); this measure captures the consistency of the response across repeats at multiple time scales. At low contrast, the coherence of the excitatory current matched that of the LN model prediction, whereas at high contrast the coherence of the current extended to finer time scales (i.e., higher frequencies) and hence exceeded the precision predicted by the LN model (Fig. 1F).

Contrast adaptation was measured in the synaptic currents by comparing LN models at each contrast level (Fig. 1E). The linear filter for the current responses had a larger amplitude (i.e., higher gain) in low contrast compared with high contrast (Beaudoin et al., 2007; Beaudoin et al., 2008; Kim and Rieke, 2001). The adaptation occurred rapidly after the contrast switch and lacked an additional slow component described for some other ganglion cell types (Suppl. Fig. 1; (Baccus and Meister, 2002; Manookin and Demb, 2006)). Furthermore, the increase in gain for currents was smaller than that measured in the spike response. To compare the changes in the filters with contrast and spikes, we define contrast gain as the ratio between the standard deviation of the filter in low contrast over that in high contrast. For all neurons where spikes and currents were recorded in the same neuron, the contrast gain was significantly larger for spikes than currents (*n*=3), and this trend was robust across all recordings (p<10^−6^, *unpaired two-sample t-test*, spikes: *n*=11, current: *n*=13) (Fig. 1E) (Zaghloul et al., 2005). These observations suggest that both contrast adaptation and temporal precision in ON-Alpha ganglion cell spike responses are generated in large part by retinal circuitry upstream of the ganglion cell, but that further transformation occurs between currents and spikes (Kim and Rieke, 2001).

**Figure 1.**
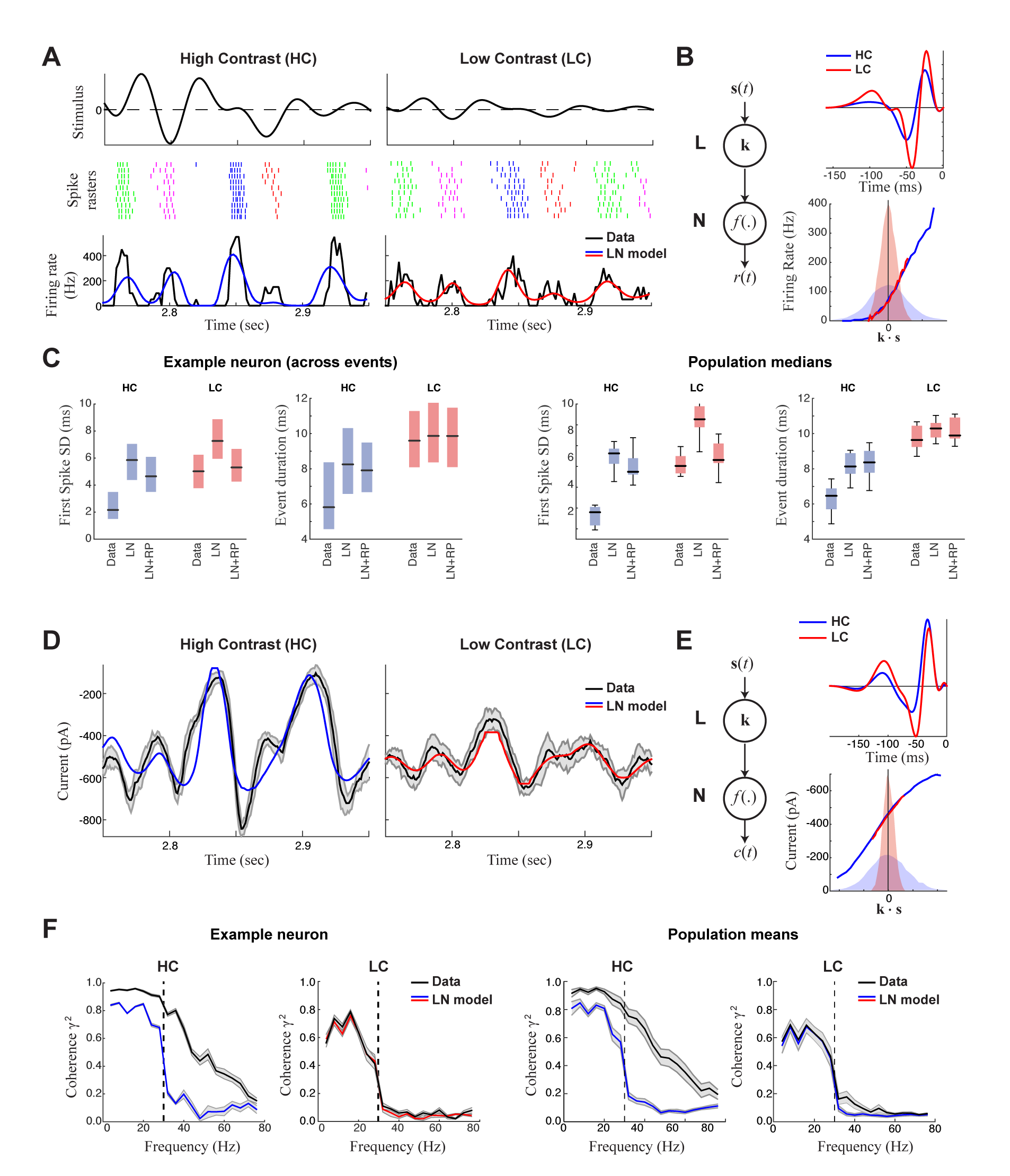
Precision of ganglion cell spike trains arises from synaptic inputs. **A.** Spike rasters of an ON-Alpha cell to 10 repeated presentations of a temporally modulated noise stimulus *(top)* at two contrast levels. The response was parsed into separate “events” (labeled by different colors). The PSTH *(bottom)* is compared with predictions of the LN model (blue, red), which fits relatively better at low contrast. **B**. The LN model (schematic: *left)* was fit separately at each contrast, with the effects of adaptation isolated to the linear filters (top), which share the same nonlinearity (bottom). Shaded distributions show the filtered stimulus at high (blue) and low (red) contrasts. **C**. Temporal properties of the observed spike trains, compared with predictions of the LN model without or with a spike-history term (LN and LN+RP). *Left:* SD of the timing of the first spike in each event. *Right:* Event duration, measured by the SD of all spikes in the event (*p<10^−6^, 59 events). LN and LN+RP models do not reproduce the spike precision at high contrast (HC), but the LN+RP model is adequate at low contrast (LC). **D**. Excitatory synaptic current from the neuron in A-C compared with the LN model predictions. Gray area indicates SD across trials, demonstrating minimal variability. **E**. LN model fits to the current data. The temporal filters *(top)* change less with contrast compared to spike filters (Fig. 1B). **F**. The precision of the current response was measured using the coherence between the response on individual trials and either the observed trial-averaged response (black) or LN predictions (blue, red). Gray area shows SEM across trials *(left)* and standard deviation across the population *(right)*. The LN model fails to capture high frequency response components at HC, but agrees well with the data at LC, suggesting the precision observed in ganglion cell spike trains arises at the level of synaptic inputs.

### The nonlinear computation underlying synaptic inputs to ganglion cells

In constructing a nonlinear description of the computation present in excitatory synaptic currents, we sought to emulate elements of the retinal circuit that shape these currents (Fig. 2A). Excitatory synaptic inputs to ganglion cells come from bipolar cells, and bipolar cell voltage responses to our stimuli are well described by an LN model (Baccus and Meister, 2002; Rieke, 2001). This suggests that mechanisms responsible for the nonlinear behavior of the postsynaptic excitatory current are localized to the bipolar-ganglion cell synapses. Possible sources of such nonlinear behavior include presynaptic inhibition from amacrine cells, which can directly gate glutamate release from bipolar cell terminals (Eggers and Lukasiewicz, 2011; Euler et al., 2014; Schubert et al., 2008; Zaghloul et al., 2007), and synaptic depression at bipolar terminals caused by vesicle depletion (Jarsky et al., 2011; Markram et al., 1998; Ozuysal and Baccus, 2012).

To capture the computations that could be performed by such suppressive mechanisms, we constructed a “divisive suppression” (DivS) model (Fig. 2A, *bottom left)*. Terms simulating bipolar cell excitation and suppression are each described by a separate LN model, with a multiplicative interaction between their outputs such that the suppression impacts bipolar cell release (Fig. 2A). Note that divisive gain control matches earlier models of either presynaptic inhibition (Olsen and Wilson, 2008) or synaptic depression (Markram et al., 1998). The suppressive term drops below one when the stimulus matches the suppressive filter, causing a proportional decrease in excitation of the ganglion cell. If the suppression does not contribute to the response, its nonlinearity would simply maintain a value of one, and the DivS model reduces to the LN model. The DivS model construction can be tractably fit to data using recent advances in statistical modeling (Ahrens et al., 2008b; McFarland et al., 2013).

**Figure 2.**
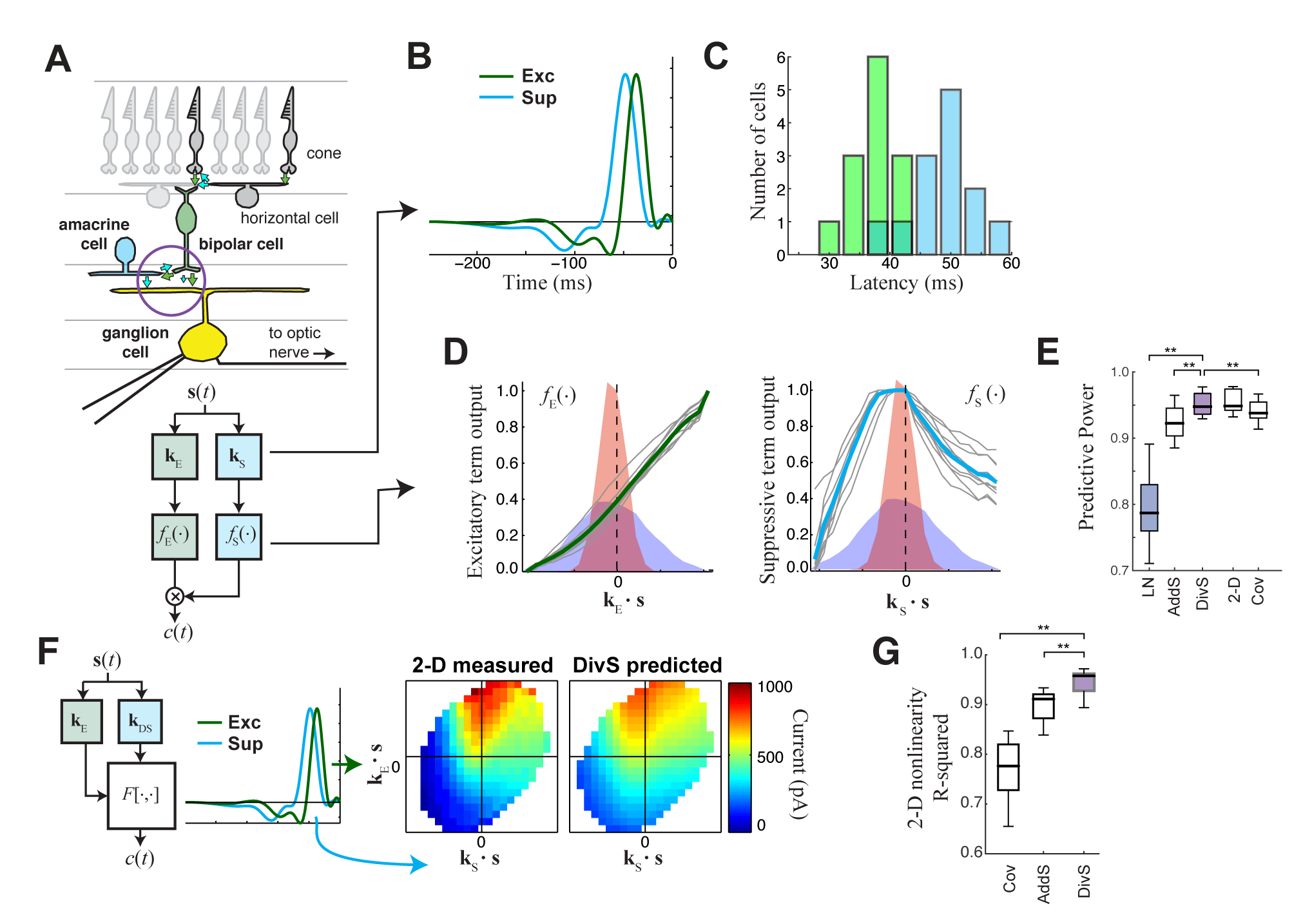
The divisive suppression (DivS) model of synaptic currents. **A.** Schematic of retinal circuitry. The vertical, excitatory pathway, cones bipolar cells ganglion cell can be modulated at the bipolar cell synapse by amacrine cell-mediated inhibition of bipolar cell release or by synaptic depression. We model both processes by divisive suppression *(bottom inset)*, where an LN model, representing the collective influence of amacrine cell inhibition and synaptic depression, multiplicatively modulates excitatory inputs from the bipolar cell. **B**. The excitatory (green) and suppressive (cyan) temporal filters of the DivS model for an example ON-Alpha cell. **C**. Divisive suppression is delayed relative to excitation, demonstrated by the distributions of latencies measured for each pair of filters (mean delay = 10.9±2.2 ms, p<0.0005, *n*=13). **D**. Excitatory *(left)* and suppressive nonlinearities *(right)* for the DivS model. The solid line indicates model fits for the example cell, and the gray lines are from other cells in the population, demonstrating their consistent form. The distribution of the filtered stimulus is also shown as the shaded area for HC (blue) and LC (red). The suppressive nonlinearity (*right*) falls below one for stimuli that match the kernel or are opposite, implying that divisive suppression is ON-OFF. **E**. To validate the form of the DivS model, we compared its performance to alternative models, including a more general model where the form of the nonlinearity is not assumed (2-D, see below), a covariance (COV) model similar to spike triggered covariance (Suppl. Fig. 2), and a model where excitatory and suppressive terms interact additively (AddS) instead of divisively. The DivS model performed significantly better than the LN, AddS and COV models (**p<0.0005, *n*=13), and matched the performance of the 2-D model. **F**. We used a 2-dimensional nonlinearity to capture any more general interaction between excitatory and suppressive filters, shown with schematic *(left)*, and the resulting fits *(middle)*. Consistent with the model performance (E), the form of this 2-D nonlinearity could be reproduced by the DivS model (*right*). **G**. Accuracy of 2-D nonlinearity reconstruction with DivS model and AddS model across neurons (\*\**p*<0.0005, *n*=13).

The DivS model fits were highly consistent across the population, with similarly shaped excitatory and suppressive filters across cells (Fig. 2B). For each cell, the suppressive filter was delayed relative to the excitatory filter (10.9 ± 2.2 ms, p<0.0005, *n*=13, Fig. 2C). The excitatory nonlinearity was approximately linear over the range of stimuli (Fig. 2D, *left*), whereas the suppressive nonlinearity decreased below one when the stimulus either matched or was opposite to the suppressive filter (Fig. 2D, *right)*, resulting in selectivity to both light increments and decrements.

The DivS model outperformed the LN model in predicting the observed currents (Fig. 2E). Furthermore, it performed as well or better than other nonlinear interactions between the two filters. We first tested a more general form of nonlinear interaction by directly estimating a two-dimensional nonlinear function, which maps each combination of the excitatory and suppressive filter outputs to a predicted current (Fig. 2F; see Methods). While this 2-D model contains many more parameters than the DivS model, the 2-D model did not perform significantly better (Fig. 2E); indeed, the estimated 2-D nonlinearities for each neuron were well approximated by the separable mathematical form of the DivS model (*R*^2^ for 2-D nonlinearity reconstruction = 0.94±0.02; Fig. 2G). We also tested an additive suppression (AddS) model, where suppression interacts with excitation additively (see Methods). However, the AddS model had significantly worse predictive power than the DivS model (p<0.0005, *n*=13; Fig. 2E) and less resemblance to the 2-D nonlinearities compared to the DivS model (p<0.0005, *n*=13; Fig. 2G). Therefore, the DivS model gives a parsimonious description of the nonlinear computation at the bipolar-ganglion cell synapse and yields interpretable model components, suggesting an interaction between tuned excitatory and suppressive elements.

We also derived filters using a form of spike-triggered covariance (Fairhall et al., 2006; Liu and Gollisch, 2015; Samengo and Gollisch, 2013) adapted for the continuous nature of the synaptic currents (see Methods). Consistent with previous analyses with spikes (Butts et al., 2011; McFarland et al., 2013), these covariance methods identified the same filter subspace as the DivS model, meaning that the covariance-based filters could be derived as a linear combination of the DivS filters and vice versa (Suppl. Fig. 2-1). However, the 2-D mapping between STC filter output and the synaptic current differed substantially from the same mapping for the DivS model (Suppl. Fig. 2-1). As a consequence, the 2-D mapping for the STC analysis could not be decomposed into two 1-D components (Suppl. Fig. 2-1). Thus, despite the ability of covariance analysis to nearly match the DivS model in terms of model performance (Fig. 2E), it could not uncover the divisive interaction between excitation and suppression (Fig. 2G). As we demonstrate below, the correspondingly straightforward divisive interaction detected by the DivS model on the ganglion cell synaptic input is essential in deriving the accurate model of ganglion cell output, which combines this divisive interaction with subsequent nonlinear components related to spike generation.

### Divisive suppression explains contrast adaptation in synaptic currents

In addition to nearly perfect predictions of excitatory current at high contrast (Fig. 2; Fig. 3C), the DivS model also predicted the slower time course of the synaptic currents at low contrast. Indeed, using a single set of parameters the model was similarly accurate in both contrast conditions (Fig. 3A), and outperformed an LN model that used separate filters fit to each contrast level (e.g., Fig. 1E). The DivS model thus implicitly adapts to contrast with no associated changes in parameters.

The adaptation of the DivS model arises from the scaling of the divisive term with contrast. The fine temporal features in the synaptic currents observed at high contrast (Fig. 3C, *left)* arise from the product of the output of the excitatory LN component and the output of the suppressive LN component.Because suppression is delayed relative to excitation and has both ON and OFF selectivity, suppression increases at both positive and negative peaks of the suppressive filter output (Fig. 3C *inset)*. This divisive suppression makes the DivS model output more transient compared to its excitatory component output alone; the difference between the two predictions is pronounced surrounding the times of peak excitation. At low contrast (Fig. 3C, *right)*, both excitatory and suppressive filter outputs are proportionately scaled down. Because the suppression is divisive and close to one, the DivS model becomes dominated by the excitatory term and closely matches the LN model, as well as the measured excitatory current.

The close match between data and DivS predictions across contrasts suggest that the DivS model should exhibit contrast-dependent change in the LN model filters (e.g., Fig. 1E). Indeed, using the LN model to describe the filtering properties of the DivS-model-predicted currents in high and low contrast matches the LN-based adaptation measured from data on a cell-by-cell basis (Fig. 3D), including the observed contrast gain (Fig. 3E) and change in biphasic index (Fig. 3F).

**Figure 3.**
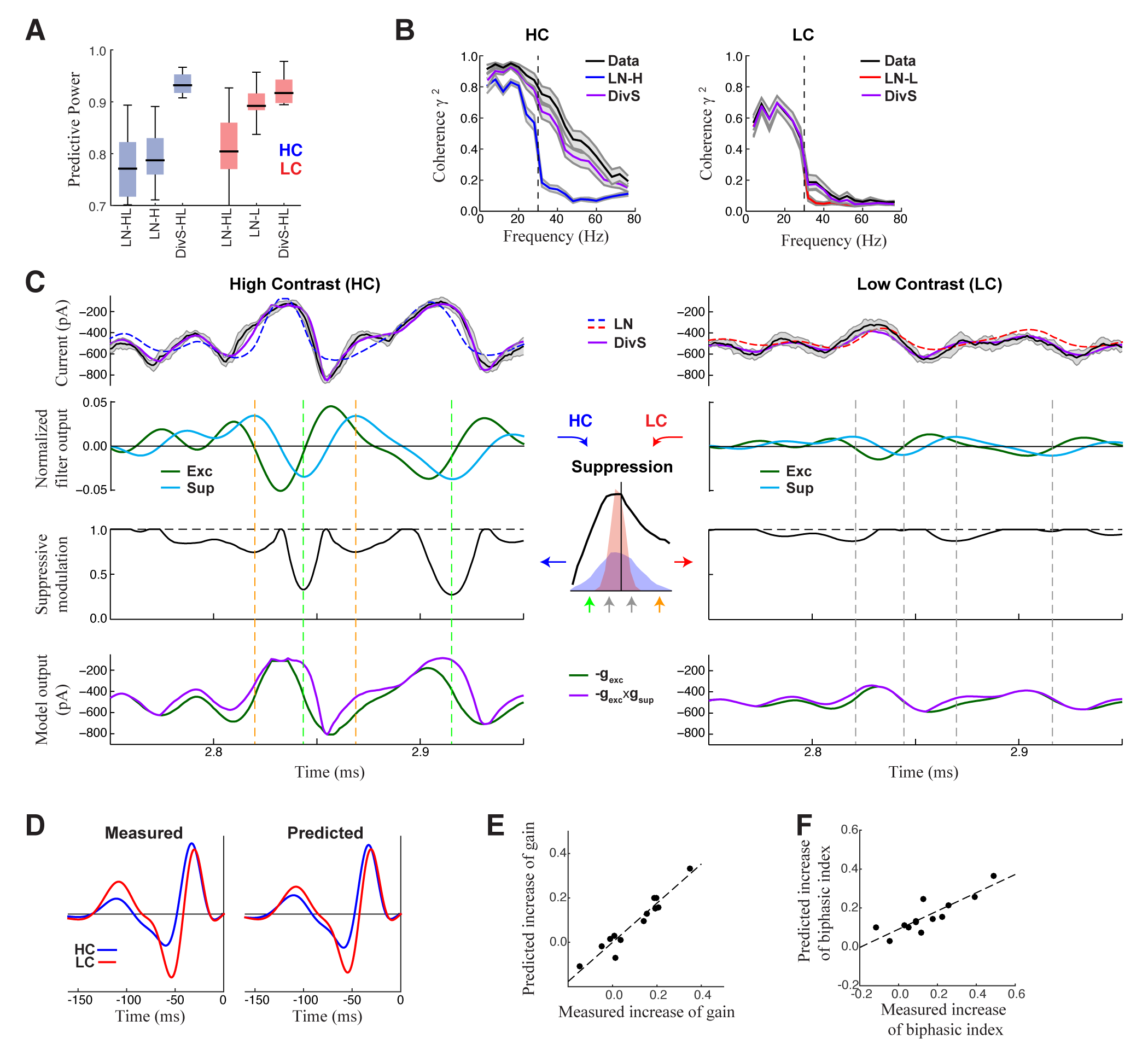
DivS model explains temporal precision and contrast adaptation in synaptic currents. **A**. The predictive power of models across contrasts. The DivS model is fit to both contrasts using a single set of parameters, and outperforms LN models fit separately to either high or low contrast (LN-H and LN-L). As expected, the LN model fit to both contrasts (LN-HL) performs worse than separately fit LN models, because it cannot capture the filter changes without changes in model parameters. **B**. Average coherence between model predictions and recorded synaptic currents on individual trials (*n*=13), shown for high contrast (HC) and low contrast (LC). The DivS model prediction is almost identical to that of the trial-averaged response. **C**. DivS model explains precision and contrast adaptation through the interplay of excitation and suppression. *Top:* comparison of predictions of synaptic current response of the LN model and the DivS model for the cell in Fig. 1. *2^nd^ row:* normalized output of the excitatory and delayed suppressive filter. *3^rd^ row:* suppressive modulation is obtained by passing the filtered output through the suppressive nonlinearity *(middle inset). Bottom:* excitatory output of the DivS model before and after the suppressive modulation. In LC, the suppressive term (*3^rd^ row*) does not deviate much from unity, and consequently the DivS model output resembles the excitatory input. **D**. Comparison of the measured *(left)* and DivS model predicted *(right)* filters across contrasts. **E-F**. The DivS model accurately captures changes of both contrast gain (E: R=0.96, p<10^−6^) and biphasic index (F: R=0.86, p<0.0005) of the temporal filters across contrasts.

### Divisive suppression largely originates from the surround region of the receptive field

As described above (Fig. 2A), the mathematical form of the DivS model is consistent with two presynaptic mechanisms that shape temporal processing: synaptic depression (Jarsky et al., 2011; Ozuysal and Baccus, 2012) and presynaptic inhibition (Eggers and Lukasiewicz, 2011; Schubert et al., 2008). Indeed, a model of ganglion cells that explicitly implements synaptic depression, the linear-nonlinear-kinetic model (LNK model) (Ozuysal and Baccus, 2012) can explain intracellular recordings. The LNK model fits a single LN filter (analogous to the excitatory **k***_E_* and *f_E_*(.) of the DivS model; Fig. 2A), with additional terms that simulate use-dependent depletion of output (Suppl. Fig. 3). This depletion is based on previous output of the model (recovering over one or more time scales), and divisively modulates the output of the LN filter. For our data, the LNK model captured excitatory currents in response to the temporally modulated spot (Fig. 4A), also outperforming the LN model (*p*<0.0005, *n* = 13), although not with the level of performance as the DivS model (*p*<0.0005, *n* = 13). Furthermore, when data were generated *de novo* by an LNK model simulation, the resulting DivS model fit showed a delayed suppressive term, whose output well approximates the effect of synaptic depression in the LNK model (Suppl. Fig. 3).

The DivS and LNK model, however, yield distinct model predictions to a more complex stimulus where a central spot and surrounding annulus are independently modulated (Fig. 4B). The models described above can be trivially extended to this stimulus by including two temporal filters, one for the center and one for the surround (Fig. 4B). As expected from the center-surround structure of ganglion cell receptive fields, an LN model fit to this condition demonstrates strong ON-excitation from the center, and a weaker OFF component from the surround.

Because the LNK model has a single filter that explains both the excitation and resulting synaptic depression, the LNK model’s filter resembled the LN filter (Fig. 4B). Furthermore, the “spatial” LNK model had rate constants that differed significantly from those fit to the single temporally modulated spot (Fig. 4C), which minimized the time that the model dwelled in the inactivated state (i.e., was “suppressed”). Correspondingly, the LNK model in the spot-annulus condition exhibited little performance improvement over the LN model (predictive power improvement 1.8% ± 1.3%, *p*=0.016, *n*=7; Fig. 4D).

By comparison, the DivS model significantly outperformed the LN model with an improvement of 8.6% ± 3.3% (*p*=0.016; *n*=7), and was 6.7 ± 2.8% better than the LNK model (*p*=0.016; *n*=7). The suppressive term of the DivS model showed a very distinct spatial profile relative to excitation, with a greater drive from the annulus region, while excitation was mostly driven by the spot region (Fig. 4E,F). The suppressive filter overlapping the spot was typically slower than the filter overlapping the annulus: the peak latency for the suppressive filter was 129 ± 16 ms within the spot region compared to 120 ± 15 ms within the annulus region (faster by 9.7 ± 4.3 ms; *p* =0.0156, *n*=7).

The strong suppression in the surround detected by the DivS model could not be explained by the LNK model, which cannot flexibly fit an explicit suppressive filter. Indeed, suppression in the LNK model arises from excitation, and thus the two components share the same spatial profile (Fig. 4B; Suppl. Figs. 3 and 4). This can be demonstrated not only with simulations of the LNK model, but also more complex models with separate LNK terms in center and surround (Suppl. Fig. 4). In all cases, application of the DivS model to data generated by these synaptic-depression-based simulations found that the suppressive term roughly matched the spatial profile of excitation, which is inconsistent with the observed data (Fig. 4F). While these analyses do not eliminate the possibility that synaptic depression plays a role in shaping the ganglion cell response (and contributes to the DivS suppression), the surround suppression suggests that synaptic depression alone cannot fully describe our results. In contrast, the DivS model can flexibly capture the more general suppression profiles of other more complex models (Suppl. Fig. 4) as well as the data (Fig. 4F), which could ultimately be related to mechanisms of synaptic depression and other sources such as pre-synaptic inhibition.

**Figure 4.**
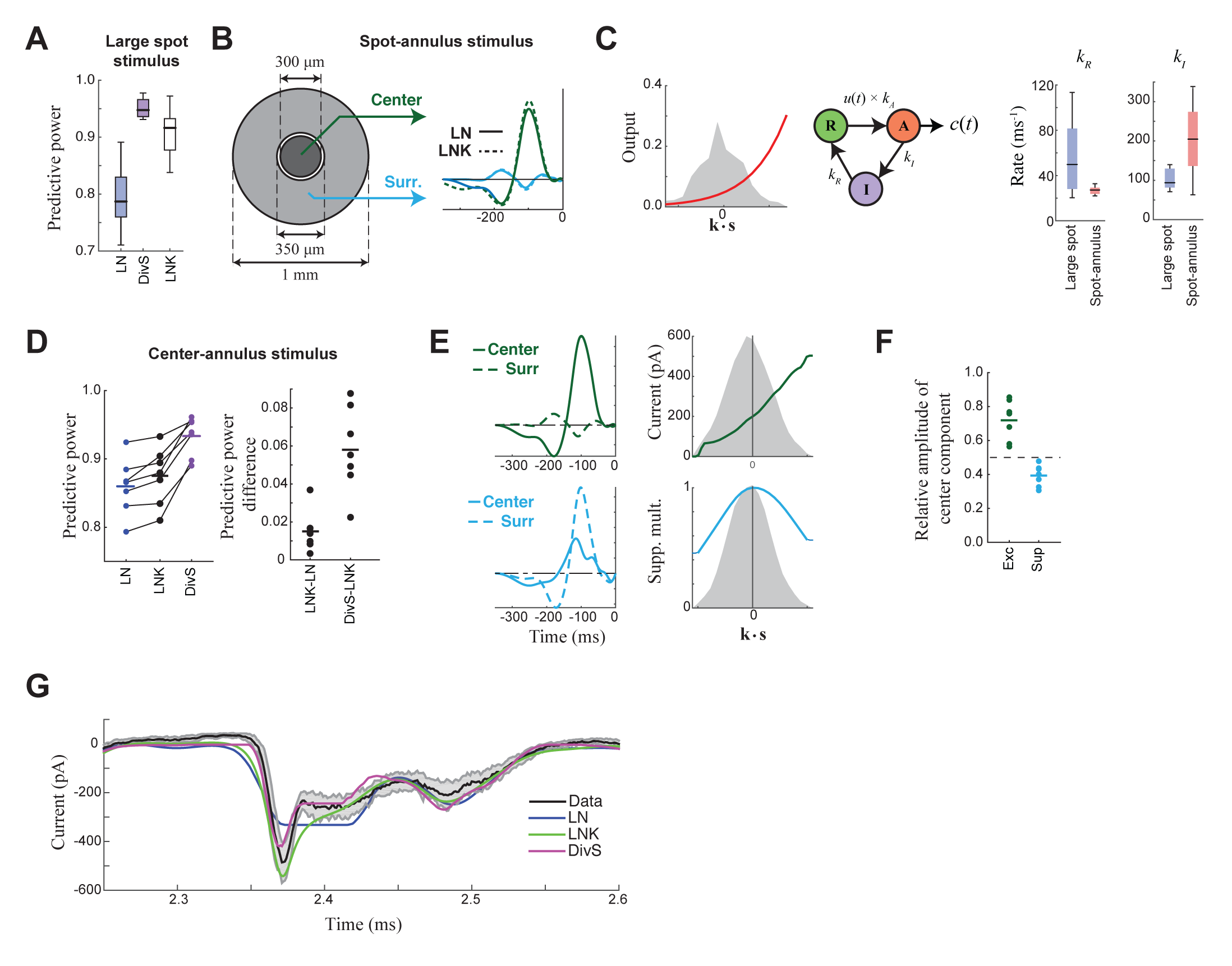
Probing the mechanism of divisive suppression with center-surround stimuli. **A**. For the large spot stimulus, the Linear-Nonlinear-Kinetic (LNK) model nearly matches the performance of the DivS model, and outperforms the LN model. **B**. To distinguish between different sources of divisive suppression, we presented a spot-annulus stimulus *(left)*, where each region is independently modulated. Model filters can be extended to this stimulus using a separate temporal kernel for center and surround, shown for the LN and LNK model filters *(right)*, which are very similar. **C.** After the linear filter, the LNK model involves a nonlinearity *(left)*, which together drive the transition between resting and activated states *(middle)*, which is further governed by kinetics parameters as shown. Critical kinetics parameters for LNK models differed between the large-spot and spot-annulus stimulus (*right*), however, with the spot-annulus model very quickly transitioning from Inactive back to Active states, minimizing the effects of synaptic depression. **D**. The performance of the spatiotemporal LNK model are only slightly better than those of the LN model, and neither captures the details of the modulation in synaptic current, compared with the DivS model. **E.** The spatiotemporal DivS model shown for an example neuron exhibits different spatial footprints for excitation and suppression, with excitation largely driven by the spot and suppression by the annulus. This divisive suppression is inconsistent with synaptic depression, which predicts overlapping sources of suppression and excitation (see Suppl. Figs. 3 and 4). **F**. Contribution of the center component in the DivS model for excitation *(left)* and suppression *(right)*. Excitation is stronger in the center than in the surround (center contribution>0.5, *p*=0.016, *n*=7) and suppression is weaker in the center (center contribution<0.5, *p*=0.016, *n*=7) for every neuron. **G.** The DivS model is able to capture temporal transients in the current response to spot-annulus stimuli better than the LN and LNK models.

### Nonlinear mechanisms underlying spike outputs of ganglion cell

With an accurate model for excitatory synaptic currents established, we returned to modeling the spike output of ON-Alpha cells. Following previous likelihood-based models of ganglion cell spikes, we added a spike-history term, which implements absolute and relative refractory periods (Butts et al., 2011; McFarland et al., 2013; Paninski, 2004; Pillow et al., 2005). The output of this spike-history term sums with the contributions of the DivS model for the synaptic currents and is further processed by a spiking nonlinearity (Fig. 5A), yielding the final predicted firing rate. Using a standard likelihood-based framework, all terms of the model – including the excitatory and suppressive LN models that comprised the prediction of synaptic currents – can then be tractably fit using spike data alone. But it is important to note that this model architecture was only made clear via the analyses of synaptic currents described above.

When fit using spiking data alone, the resulting excitatory and suppressive filters and nonlinearities closely resembled those found when fitting the model to the synaptic currents recorded from the same neurons (e.g., Fig. 2B,D). Suppression was consistently delayed relative to excitation (Fig. 5B), and exhibited both ON and OFF selectivity (Fig. 5C). The spike-history term was suppressive and had two distinct components, a strong “absolute refractory period” that lasted 1-2 ms and a second relative refractory period that lasted more than 15 ms (Berry and Meister, 1998; Butts et al., 2011; Keat et al., 2001; Paninski, 2004; Pillow et al., 2005).

The resulting model successfully captured over 90% of the predictable variance in the firing rate for all neurons in the study (Fig. 5F, median=91.5% ± 1.0%; *n*=11), representing the best model performance for ganglion cell spike trains considered at millisecond resolution. By comparison, the standard LN model had a median predictive power of 62.8% ± 1.9% (*n* = 11); which modestly increased to 68.8% ± 1.9% upon inclusion of a spike-history term (Fig. 5F). This suggests that ganglion cell spikes are strongly shaped by the nonlinear computations present at their synaptic input, and the precise timing of ganglion cell spiking involved the interplay of divisive suppression in their input with spike-generating mechanisms.

**Figure 5.**
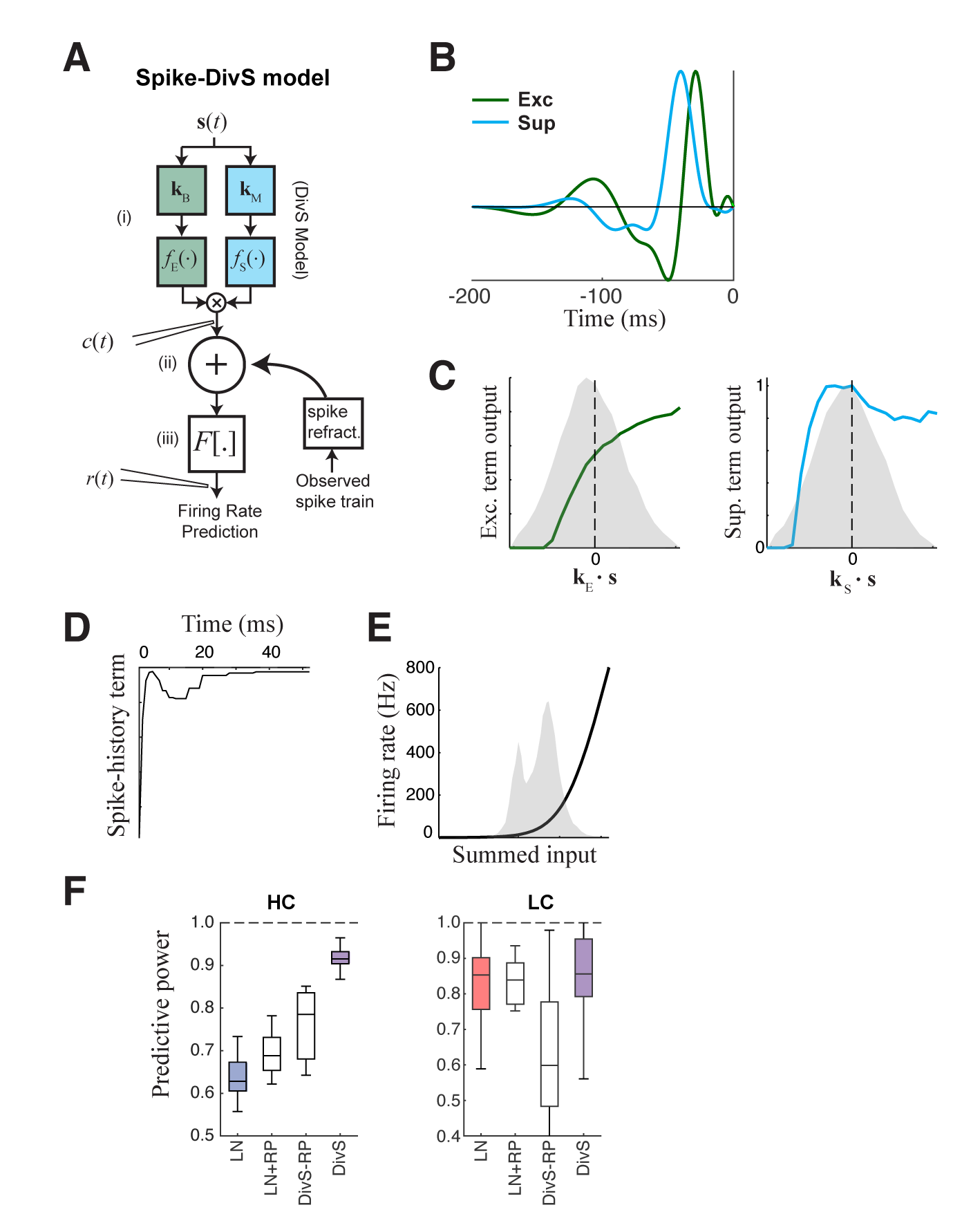
The extended divisive suppression model explains ganglion cell spike trains with high precision. **A**. Model schematic for the divisive suppression model of spiking, which extends DivS model for the current data by adding an additional suppressive term for spike-history (refractoriness), with the resulting sum passed through a rectifying spiking nonlinearity. (B-E) The model components for the same example neuron considered in Fig. 1-3. **B**. The excitatory and suppressive filters. **C**. The excitatory and suppressive nonlinearities. The filters and nonlinearities are similar to the DivS model fit from current data (shown in Fig. 2B). **D**. The spike-history term, demonstrating an absolute and relative refractory period. **E**. The spiking nonlinearities, with shaded area indicating the distribution of generating signals. **F**. The predictive power of different models applied to the spike data in HC and LC. The DivS model performs better than other models (HC: *p*<0.001; LC: *p*<0.002, *n*=11), including the LN model, the LN model with spike history term (LN+RP), and a divisive suppression model lacking spike refractoriness (DivS-RP). Only a single set of parameters was used to fit the DivS model for both contrasts, whereas all other models shown used different parameters fit to each contrast.

### Precision of spike trains arises from complementary mechanisms of divisive suppression and spike refractoriness

To evaluate the relative contributions of divisive suppression and spike refractoriness to predicting firing, we simulated spike trains using different combinations of model components (Fig. 6A). We first found that the parameters of the divisive suppression components could not be fit without including a spike-history term, suggesting that each component predicts complementary forms of suppression. We could generate a DivS model without a spike-history term, however, by first determining the full model (with spike-history term), and then removing the spike history term and refitting (see Methods), resulting in the DivS-RP model. This allowed for direct comparisons between models with selective deletion of either divisive suppression or spike refractoriness (Fig. 6A).

Event analyses on the resulting simulated spike trains, compared with the observed data, demonstrate that both divisive suppression (derived from the current analyses above) and spike refractoriness were necessary to explain the precision and reliability of ganglion cell spike trains. By comparing the two models without DivS (LN and LN+RP) to those with DivS (DivS and DivS-RP), it is clear that divisive suppression is necessary to predict the correct envelope of the firing rate (Fig. 6B). Note, however, that DivS had little impact in the low contrast condition, which lacked fine-time-scale features of the spike response.

By comparison, the spike-history term had little effect on the envelope of firing (Fig. 6A, *bottom*), and contributed little to the fine time scales in the ganglion cell spike train at high contrast (Fig. 6B). Instead, the spike-history term had the largest effect on correct predictions of event reliability, as reflected in the event Fano factor (Fig. 6C). Both models without the spike-history term had much variability in spike counts in each event. The presence of the suppression contributed by the spike-history term following each event allows the predicted firing rate to be much higher (and more reliable) during a given event, resulting in reliable patterns of firing within each event (Fig. 6A) (Pillow et al., 2005).

We conclude that a two-stage computation present in the spike-DivS model, with both divisive suppression and spike refractoriness, is necessary to explain the detailed spike patterning on ON-Alpha ganglion cells.

**Figure 6.**
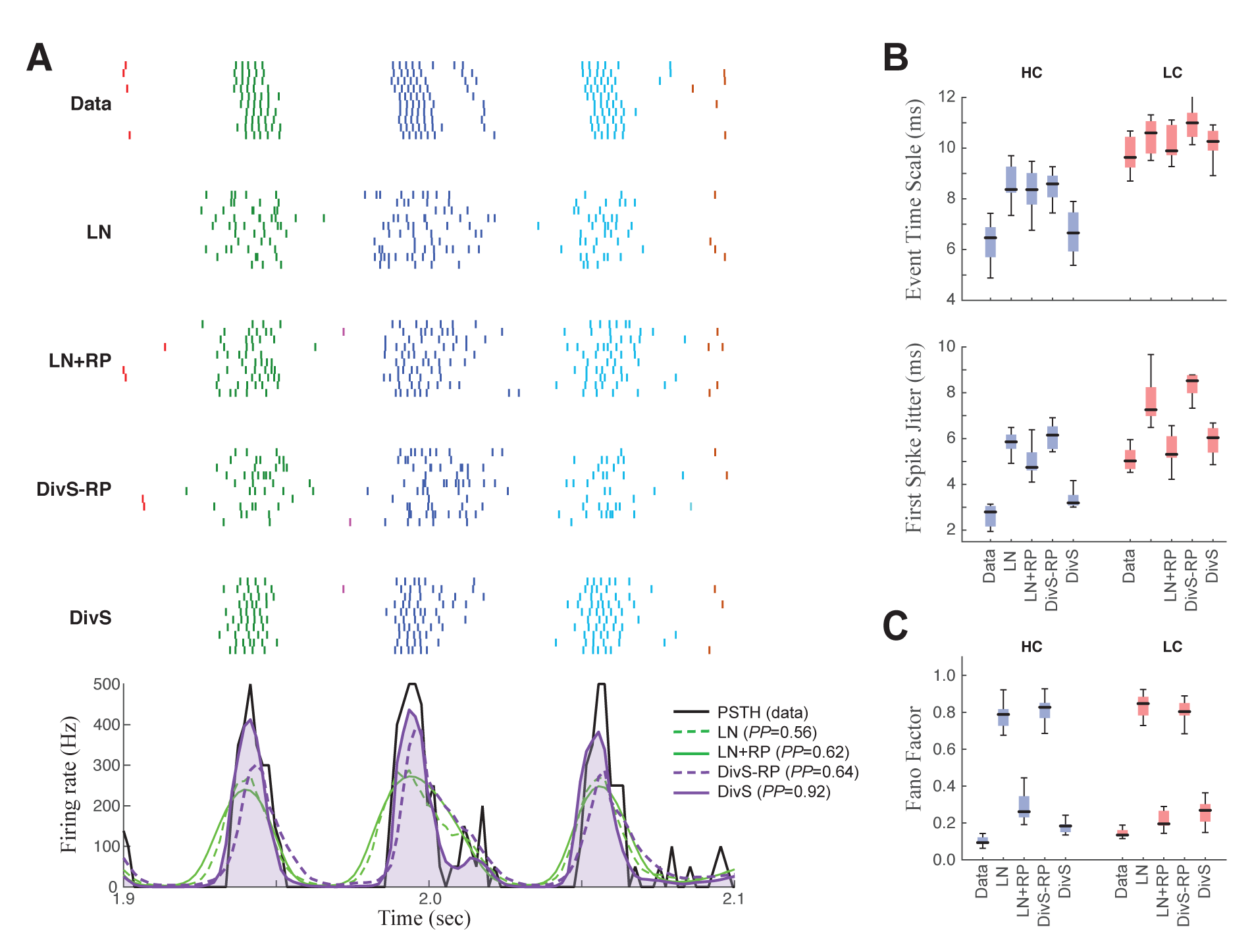
Spike patterning is shaped by a combination of nonlinear mechanisms. **A.** *Top*: Spike rasters recorded over ten repeats for an example cell (black) compared with simulated spikes from four models: LN, LN model with spike-history term (LN+RP), the DivS model without spike-history (DivS-RP), and the full DivS model (DivS). Colors in the raster label separate spike events across trials (see Methods). *Bottom*: The PSTHs for each model demonstrate that suppressive terms are important in shaping the envelope of firing (DivS prediction is shaded). (**B-E**) Using event labels, spike statistics across repeats were compiled to gauge the impact of different model components. **B**: The temporal properties of events compared with model predictions, across contrast (same as Fig. 1, with DivS-based models added). Both spike-history and divisive suppression contribute to reproduce the temporal scales across contrast. **C.** The Fano factor for each event is a measure of reliability, which increased (i.e., Fano factor decreased) for models with a spike-history term.

### Contrast adaptation is enhanced via spike refractoriness in ganglion cell output

In addition to accurate reproduction of precise spike outputs of ganglion cells, the DivS model also captured the effects of contrast adaptation observed in the ganglion cell spike trains. For both contrast conditions, the simulated spike trains, which are predicted for both contrasts using a single set of parameters, are almost indistinguishable from the data (Fig. 7A, *top*). As with the performance of the models of excitatory current (Fig. 3), the DivS model outperforms LN models that are separately fit for each contrast level (Fig. 5F).

The ability to correctly predict the effects of contrast adaptation depends on both the divisive suppression and spike-refractoriness of the spike-DivS model. This can be shown for an example neuron by using LN filters of the simulated output of each model at high and low contrasts (Fig. 7B). In this case, only the DivS model (which includes a spike-history term) shows adaptation similar to that observed by the LN filters fit to the data. We quantified this across the population by identifying the most prominent feature of adaptation of the LN filters, the change in filter amplitude (i.e., contrast gain). Across the population, the DivS correctly predicted the magnitude of this change (Fig. 7C, *top*), as well as the changes in biphasic index across contrasts (Fig. 7C, *bottom*), and outperformed models with either the divisive suppression or spike-history terms missing.

As expected, spike refractoriness imparted by the spike-history term contributed to the stronger effects of contrast adaptation observed in spikes relative to synaptic inputs (Beaudoin et al., 2007; Kim and Rieke, 2001, 2003; Rieke, 2001; Zaghloul et al., 2005). Specifically, at high contrast, spikes concentrate into relatively smaller time windows, leading to a consistently timed effect of spike refractoriness (Fig. 7D). As a result, despite similar numbers of spikes at the two contrasts, the effect of the spike-history term has a bigger impact at high contrast.

Thus, contrast adaptation – and more generally the temporal shaping of ON-Alpha ganglion cell spike trains – depends on nonlinear mechanisms at two stages of processing within the retinal circuit: with precision emerging predominantly from ganglion cell synaptic inputs, and spike-refractoriness amplifying the effects of precision to explain spike patterning and adaptation to contrast.

**Figure 7.**
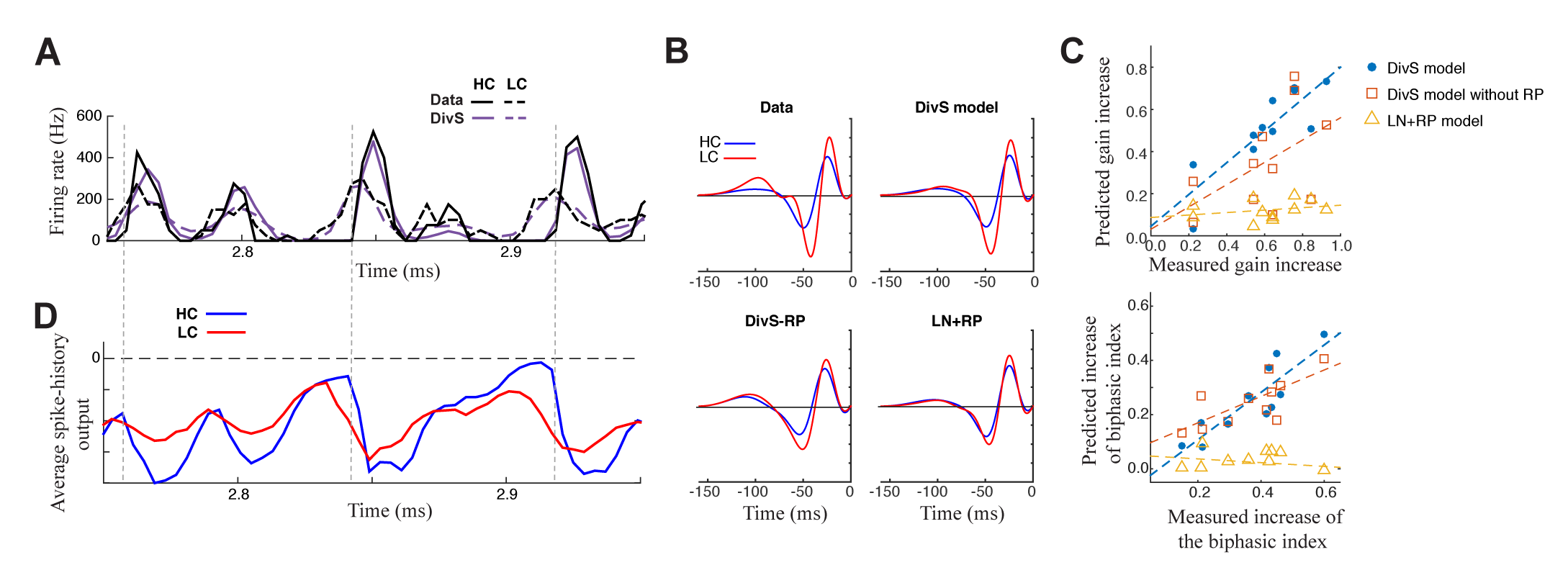
Contrast adaptation in the spike output depends on both divisive suppression and spike refractoriness. **A.** The full spike-DivS model accurately captured contrast adaptation. *Top:* observed PSTH and predicted firing rates of the DivS model at HC and LC. **B.** The DivS model predicts the changes in LN filter shape and magnitude with contrast for an example cell. Predicted changes are shown for each model, demonstrating that the full effects of contrast adaptation require both divisive suppression and spike-history terms. **C**. Measured and predicted contrast gain (*top*) and changes of biphasic index (*bottom*). The DivS model accurately predicts contrast gain and changes biphasic index across contrast across cells (contrast gain: slope of regression=0.75, *R*=0.85, *p*<0.001; biphasic index: slope of regression=0.87, *R*=0.87, *p*<0.001). DivS model without the spike history term underestimated contrast adaptation (contrast gain: slope of regression=0.53, *R*=0.51, *p*=0.10; biphasic index: slope of regression=0.49, *R*=0.73, *p*<0.05), and the LN+RP model failed to predict adaptation altogether (contrast gain: slope of regression=0.06, *R*=0.28, *p*=0.41; biphasic index: slope of regression=−0.07, *R*=−0.18, *p*=0.60). **D**. The suppressive effect from the spike-history term is amplified at HC, due to increased precision of the spike train. Dashed lines show the onset of HC spike events, which predict the largest difference in magnitudes of the suppression between contrasts.

## DISCUSSION

In this study we derived a retina-circuit-inspired model for ganglion cell computation using recordings of both the synaptic inputs and spike outputs of the ON-Alpha ganglion cell. Data were used to fit model parameters and evaluate different hypotheses of how the retinal circuit processed visual stimuli. The resulting model explains both high precision firing and contrast adaptation with unprecedented accuracy. Precise timing is already present in the excitatory synaptic inputs, and can be explained by divisive suppression, which likely depends on a combination of mechanisms: presynaptic inhibition of bipolar terminals from amacrine cells and synaptic depression at bipolar cell synapses. The interplay between nonlinear mechanisms, including divisive suppression, spike refractoriness and spiking nonlinearity, accurately captured detailed structure in the spike response across contrast levels.

Divisive suppression was implemented by multiplying two LN models together (Fig. 2A). One LN model controls the gain of a second, simple LN model; the gain is equal to or less than one and so represents division. While divisive gain terms have been previously suggested in the retina – particularly in reference to early models of contrast adaptation (Mante et al., 2008; Meister and Berry, 1999; Shapley and Victor, 1978) – critical novel elements of the present DivS model include the ability to fit the nonlinearities of both LN terms by themselves, as well as their tractability in describing data at high time resolution. The presence of nonlinearities that are fit to data in the context of multiplicative interactions distinguishes this model from multi-linear models (two linear terms multiplying) (Ahrens et al., 2008a; Williamson et al., 2016), as well as more generalized LN models such as those associated with spike-triggered covariance (Fairhall et al., 2006; Samengo and Gollisch, 2013; Schwartz et al., 2006). Furthermore the model form allows for inclusion of spike-history terms as well as spiking nonlinearities, and can be tractably fit to both synaptic currents and spikes at high time resolution (~1 ms).

An eventual goal of our approach is to characterize the nonlinear computation performed on arbitrarily complex spatiotemporal stimuli. Here, we focused on temporal stimuli, which drive well-characterized nonlinearities in ganglion cell processing including temporal precision (Berry and Meister, 1998; Butts et al., 2007; Keat et al., 2001; Passaglia and Troy, 2004; Uzzell and Chichilnisky, 2004) and contrast adaptation (Kim and Rieke, 2001; Meister and Berry, 1999; Shapley and Victor, 1978) but do not require a large number of additional parameters to specify spatial tuning. By comparison, studies that focused on characterizing nonlinearities in spatial processing (Freeman et al., 2015; Gollisch, 2013; Schwartz and Rieke, 2011) have not modeled responses at high temporal resolution. Ultimately, it will be important to combine these two approaches, to capture nonlinear processing within spatial ‘subunits’ of the ganglion cell receptive field, and thereby predict responses at both high temporal and spatial resolutions to arbitrary stimuli. Such an approach would require a large number of model parameters and consequently a larger amount of data than collected here. Our intracellular experiments were useful for deriving model architecture – discerning the different time courses of excitation, suppression, and spike refractoriness – but ultimate tests of full spatiotemporal models will likely require prolonged, stable recordings of spike trains, perhaps using a multielectrode array.

### Generation of temporal precision in the retina

One important nonlinear response property of early sensory neurons is high temporal precision. Temporal precision of spike responses has been observed in the retinal pathway with both noise stimuli (Berry et al., 1997; Reinagel and Reid, 2000) and natural movies (Butts et al., 2007). Precise spike timing suggests a role for temporal coding in the nervous system (Berry et al., 1997), or alternatively simply suggests that analog processing in the retina must be oversampled in order to preserve information about the stimulus (Butts et al., 2007). Temporal precision also has been shown to play an important role in downstream processing of information provided by ganglion cells (Stanley et al., 2012; Usrey et al., 2000).

The generation of temporal precision involves nonlinear mechanisms within the retina; which may include both spike-refractoriness within ganglion cells (Berry and Meister, 1998; Keat et al., 2001; Pillow et al., 2005) and the interplay of excitation and inhibition (Baccus, 2007; Butts et al., 2016; Butts et al., 2011). Such distinct mechanisms contributing to ganglion cell computation are difficult to distinguish using recordings of the spike outputs alone, which naturally reflect the total effects of all upstream mechanisms. By recording at two stages of the ganglion cell processing, we demonstrate that high temporal precision already presents in the synaptic current inputs at high contrast, and temporal precision of both current inputs and spike outputs can be accurately explained by the divisive suppression model.

The divisive suppression model explained fast changes in the neural response through the interplay of excitation and suppression. For both the spike and current models, suppression is consistently delayed relative to excitation. The same suppression mechanism also likely underlies high temporal precision of LGN responses, which can be captured by a model with delayed suppression (Butts et al., 2011). Indeed, precision of LGN responses is apparently inherited from the retina and enhanced across the retinogeniculate synapse (Butts et al., 2016; Carandini et al., 2007; Casti et al., 2008; Rathbun et al., 2010; Wang et al., 2010b). Therefore, our results demonstrate that the temporal precision in the early visual system likely originates from nonlinear processing in the inputs to retinal ganglion cells. Note that the full model did not incorporate any form of direct synaptic inhibition onto the ON Alpha cell, consistent with findings that such inhibition is relatively weak and that excitation dominates the light response in the regime that we studied (Kuo et al., 2016; Murphy and Rieke, 2006).

Our results show that the contribution of spike history term to precision – as measured by the time scale of events and first-spike jitter – seems minor, consistent with earlier studies in the LGN (Butts et al., 2016; Butts et al., 2011). Nevertheless, the spike history term does play an important role in spike patterning within the event (Pillow et al., 2005) and the resulting neuronal reliability (Berry and Meister, 1998). In fact, we could not fit the divisive suppression term robustly without the spike history term in place, suggesting that both nonlinear mechanisms are important to explain ganglion cell firing.

### Contrast adaptation relies on both divisive suppression and spike refractoriness

Here we modeled contrast adaptation at the level of synaptic currents and spikes from the same ganglion cell type. We found contrast adaptation in synaptic inputs to ganglion cells, consistent with previous studies (Beaudoin et al., 2007; Kim and Rieke, 2001; Rieke, 2001; Zaghloul et al., 2005). Such adaptation could be explained by divisive suppression, which takes a mathematical form similar to previously proposed gain control models (Heeger, 1992; Shapley and Victor, 1979). Because the suppressive nonlinearity has very different shape than the excitatory nonlinearity, divisive suppression has a relatively strong effect at high contrast and results in a decrease in measured gain. Moreover, the same divisive suppression mechanism may also explain nonlinear spatial summation properties of ganglion cells (Shapley and Victor, 1979), because suppression generally has broader spatial profiles than excitation.

Contrast adaptation is amplified in the spike outputs mostly due to spike refractoriness and changes of neural precision across contrast. At high contrast, the response has higher precision and occurs within shorter event windows (Butts et al., 2010). As a result, the accumulated effect of spike refractoriness is stronger within each response event. Note that the effect of the spike history term is highly dependent on the ability of the model to predict high temporal precision at high contrast, which largely originates from the divisive suppression term as discussed earlier. Therefore, the two nonlinear properties of retinal processing, contrast adaptation and temporal precision, are tightly related mechanistically and can be simultaneously explained by the divisive suppression model.

### Circuits and mechanisms underlying the divisive suppression

Divisive suppression has been observed in many systems, including the invertebrate olfactory system (Olsen and Wilson, 2008), the lateral geniculate nucleus (Bonin et al., 2005), the primary visual cortex (Heeger, 1992), and higher visual areas like area MT (Simoncelli and Heeger, 1998). A number of biophysical and cellular mechanisms that could implement divisive suppression have been proposed, including shunting inhibition (Abbott et al., 1997; Carandini et al., 1997; Hao et al., 2009), synaptic depression (Abbott et al., 1997), presynaptic inhibition (Olsen and Wilson, 2008; Zhang et al., 2015) and fluctuation in membrane potential due to ongoing activity (Finn et al., 2007).

We evaluated different mechanistic explanations of the divisive suppression identified in this study. Divisive suppression underlying synaptic inputs to ganglion cells cannot be attributable to fluctuations in membrane potential or shunting inhibition since we recorded synaptic currents under voltage-clamp conditions that minimize inhibitory inputs. Although synaptic depression could also explain fast transient responses and contrast adaptation (Ozuysal and Baccus, 2012), this model predicts that excitation and suppression have the same spatial profiles, whereas we show that excitation and suppression have distinct spatial profiles (Fig. 4). Therefore, the divisive suppression in our model apparently depends partly on presynaptic inhibition from amacrine cells, which can extend their suppressive influence laterally (Euler et al., 2014; Schubert et al., 2008).

Detailed anatomical studies suggest that each ganglion cell type receives inputs from a unique combination of bipolar and amacrine cell types, contributing to a unique visual computation (Baden et al., 2016). By focusing on a single cell type, the ON-Alpha cell, we identified a particular computation consistent across cells. We expect that other ganglion cell types will perform different computations, and likewise have different roles in visual processing. This could include additional contrast-dependent mechanisms, including slow forms of adaptation (Baccus and Meister, 2002; Manookin and Demb, 2006), sensitization (Kastner and Baccus, 2014) and complex changes in filtering (Liu and Gollisch, 2015). Thus, further applications of the approach described here will uncover a rich diversity of computation constructed by retinal circuitry to format information for downstream visual processing.

## METHODS

### Neural recordings

Data were recorded from ON-Alpha ganglion cells from the *in vitro* mouse retina using procedures described previously (Borghuis et al., 2013; Wang et al., 2011). Spikes were recorded in the loose-patch configuration using a patch pipette filled with Ames medium, and synaptic currents were recorded using a second pipette filled with intracellular solution (in mM): 110 Cs-methanesulfonate; 5 TEA-Cl, 10 HEPES, 10 BAPTA, 3 NaCl, 2 QX-314-Cl, 4 ATP-Mg, 0.4 GTP-Na2, and 10 phosphocreatine-Tris2 (pH 7.3, 280 mOsm). Lucifer yellow was also included in the pipette solution to label the cell using a previously described protocol (Manookin et al., 2008). The targeted cell was voltage clamped at *E_Cl_* (-67 mV) to record excitatory currents after correcting for the liquid junction potential (-9 mV). Cells in the ganglion cell layer with large somas (20-25 μm diameter) were targeted. Cells were confirmed to be ON-Alpha cells based on previously established criteria (Borghuis et al., 2013): (1) an ON response; (2) high rate of spontaneous firing; and a high rate of spontaneous excitatory synaptic input; (3) a low input resistance (~40-70 MQ). In some cases, we imaged the morphology of recorded cells and confirmed (4) a relatively wide dendritic tree (300-400 μm diameter) and (5) stratification on the vitreal side of the nearby ON cholinergic (starburst) amacrine cell processes.

We made recordings from 27 ON-Alpha cells total, each in one or more of the experimental conditions described. Of the 15 cells recorded in cell-attached configuration (spike recordings), 4 cells were excluded where low reliability across trials indicated an unstable recording, as indicated by much higher spike event Fano Factors (>0.2, see below).

All procedures were conducted in accordance with National Institutes of Health guidelines under protocols approved by the Yale University Animal Care and Use Committee.

### Visual Stimulation

The temporally modulated spot stimulus was described previously (Wang et al., 2011). The retina was stimulated by UV LEDs (peak, 370 nm; NSHU-550B; Nichia America) to drive cone photoreceptors in the ventral retina. UV LEDs were diffused and windowed by an aperture in the microscope’s fluorescence port, with intensity controlled by *pClamp 9* software via a custom non-inverting voltage-to-current converter using operational amplifiers (TCA0372; ON Semiconductor). The stimulus was projected through a 4X objective lens (NA, 0.13). The stimulus was a flickering spot (1-mm diameter), with intensity generated from low pass Gaussian noise with a 30 Hz cutoff frequency. We used a contrast-switching paradigm (Baccus and Meister, 2002; Kim and Rieke, 2001; Zaghloul et al., 2005), in which the temporal contrast alternately stepped up or down every 10 sec. The contrast of the stimulus is defined by the SD of the Gaussian noise and was either 0.3 times (high contrast) or 0.1 times (low contrast) the mean. Note that this is only a three-fold difference in contrast versus the seven-fold difference considered in Ozuysal and Baccus (2012), but sufficient to see clear contrast effects. The stimulus comprised 10 cycles of 10 sec for each contrast. The first 7 sec were unique in each cycle (used for fitting model parameters), and the last 3 sec were repeated across cycles (used for cross-validation of model performance).

The center-surround stimuli (Fig. 4B) were generated in *Matlab* (Mathworks, Natick) using the *Psychophysics Toolbox* (Brainard, 1997) and presented with a video projector (M109s DLP; Dell, or identical HP Notebook Companion; HP), modified to project UV light (single LED NC4U134A, peak wavelength 385 nm; Nichia) as previously described (Borghuis et al., 2013). The center and surround stimuli were independently modulated with Gaussian noise (60-Hz update rate). A spot covered the receptive field center (e.g., 0.3 mm), and an annulus extended into the surround (e.g., inner/outer diameters of 0.35/1.0 mm). We recorded 7 ON-Alpha cells in this condition. For a subset of the recordings (*n*=5), we explored a range of inner/outer diameters, and selected the diameters that maximized the difference between the spatial footprints of excitatory and suppressive terms of the DivS model (see below).

The mean luminance of the stimulus was calculated to evoke ~4×10^4^ photoisomerizations cone^−1^ sec^−1^, under the assumption of a 1 μm^2^ cone collecting area. For all methods of stimulation, the gamma curve was corrected to linearize output, and stimuli were centered on the cell body and focused on the photoreceptors. We verified that the relatively short stimulus presentation did not result in significant bleaching, as the response (and model parameters) had no consistent trends from the beginning of the experiment to the end (Suppl. Fig. 1).

### Statistical modeling of the synaptic current response

We modeled the synaptic current response of neurons using the traditional linear-nonlinear (LN) cascade model (Paninski, 2004; Truccolo et al., 2005), as well as the Linear-Nonlinear-Kinetic model (Ozuysal and Baccus, 2012), and a general 2-D nonlinear model (‘2-D’) and Divisive Suppression model (‘DivS’) introduced in this paper.

In all cases (with the exception of the LN analyses of contrast adaptation effects described below), we optimized model parameters to minimize the mean-squared error (MSE) between the model-predicted and observed currents:

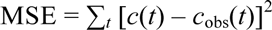

To limit the number of model parameters in the minimization of MSE, we represented temporal filters by linear coefficients weighting a family of orthonormalized basis functions (Keat et al., 2001):

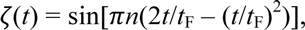

where *t*_F_=200 ms.

***LN model*.** The LN model transforms the stimulus s(*t*) to the synaptic current response *c*(*t*) using a linear filter **k**_LN_ and nonlinearity *f*_LN_[.] such that:

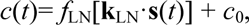

where *c*_0_ is a baseline offset. The filter **k**_LN_ was represented as a set of coefficients weighting the basis functions of eq (2), and the nonlinearities were represented as coefficients weighting tent basis functions as previously described (Ahrens et al., 2008b; McFarland et al., 2013).

***2-D model*.** We generalized the LN model by incorporating a second filtered input, such that:

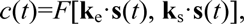

where *F*[ ·,· ] is a two-dimensional nonlinearity, and **k**_e_ and **k**_s_ denote the excitatory and suppressive filters respectively.

The 2-D nonlinearity was represented using piecewise planar surfaces and can be estimated nonparametrically for a given choice of filters (Toriello and Velma, 2012). Specifically, we divided the 2-D space into a set of uniform squares, and then subdivided each square into two triangles. Each basis function was defined as a hexagonal pyramid function centered at one of the vertices, and the 2-D nonlinearity function was expressed as a combination of these bases:

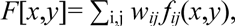

where *f_ij_*(*x*,*y*) is the basis centered at the ij_th_ grid vertex, and *w_ij_* is the weight coefficient, which can be optimized by minimizing MSE for a given choice of filters.

The coefficients for the filters and nonlinearities were optimized using block-coordinate descent: for a given choice of nonlinearity *F*[·,· ] the filters were optimized, and vice versa. In doing so, we introduced an additional constraint on the nonlinearity due to a degeneracy in the combined optimization of stimulus filters and 2-D nonlinearity. Specifically, one can choose a linear combination of the two stimulus filters and achieve the same model performance by refitting the 2-D nonlinearity. To alleviate this problem, we constrained the 2-D nonlinearity to be monotonically increasing along the first dimension, i.e.,

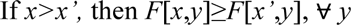

***DivS model*.** We derived the DivS model as a decomposition of the 2-D nonlinearity into two onedimensional LN models that interact multiplicatively:

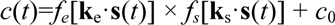

We constrained the excitatory nonlinearity *f_e_*(.) to be monotonically increasing, and constrained the second nonlinearity *f_s_*(.) to be suppressive by bounding it between zero and one, with the value for zero input constrained to be one. We optimized the filters and the nonlinearities through block-coordinate descent until a minimum MSE was found. Because this optimization problem is in general non-convex (i.e., not guaranteed to have a single global minimum), we used standard approaches (McFarland et al., 2013) such as testing a range of initialization and block-coordinate descent procedures to ensure optima were found.

***LNK Model*.** We explicitly followed the methods of Ozuysal and Baccus (2012) in fitting the linear-nonlinear-kinetic (LNK) model to the temporally modulated spot data (Fig. 4A, Suppl. Fig. 3). This model involves the estimation of an LN model in combination with a first-order kinetics model that governs the dynamics of signaling elements in resting state (R), active state (A) and inactivated states (I). The kinetics rate constants reflect how fast the signaling elements transition between states. Model parameters are fit to the data using constrained optimization. We adapted this model to fit the spot- annulus data by extending the linear filter of the LN model into separate temporal filters for center and surround processing (Fig. 4B). Note that parameters for more complex forms of the LNK model (e.g., those considered in Suppl. Fig. 4) cannot be tractably fit to real data, and we chose the parameters of these models and simulate their output, as described in Supplemental Figure 4.

### Statistical modeling of the spike response

We have applied several statistical models to describe the spike response of ganglion cells. We first considered the generalized linear modeling (GLM) framework (Paninski, 2004; Truccolo et al., 2005). We assumed that spike responses are generated by an inhomogeneous Poisson process with an instantaneous rate. The GLM makes prediction of the instantaneous firing rate of the neuron r(*t*) based on both the stimulus s(*t*) and the recent history of observed spike train **R**(*t*):

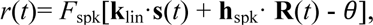

where **k**_lin_ is a linear receptive field, **h**_spk_ is the spike history term and *θ* is the spiking threshold. Here, the parameters of the model are all linear functions inside the spiking nonlinearity *F*_spk_[.]. The LN model consists of only the linear receptive field and the spiking threshold; the full GLM further includes the spike history term (denoted as LN+RP in figures). The spiking nonlinearity has a fixed functional form *F*_spk_[g] = log[1+exp(g)], satisfying conditions for efficient optimization (Paninski, 2004). The choice of this particular parametric form of spiking nonlinearity was verified with standard non-parametric estimation of the spiking nonlinearity (Fig. 1B, 1E) (Chichilnisky, 2001). The model parameters are estimated using maximum likelihood optimization. The log-likelihood (*LL*) of the model parameters that predict a firing rate *r*(*t*) given the observed neural response *r*_obs_(*t*) is (Paninski, 2004):

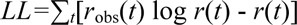

The optimal model parameters can then be determined using gradient-descent based optimization of *LL*. Although this formulation of the model assumes probabilistic generation of spikes, this fitting procedure is able to simulate and capture the parameters of the equivalent integrate-and-fire neuron, and thus makes no assumptions about the form of noise in spike generation (Butts et al., 2011; Paninski et al., 2007).

To capture nonlinear properties of the spike response, we extended the Nonlinear Input Model (NIM) (McFarland et al., 2013) to include multiplicative interactions. The predicted firing rate of the NIM is given as,

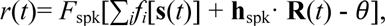

where *f*[.] represents a set of nonlinear subunits reflecting upstream processing. In this case, based on knowledge of nonlinear processing in the synaptic current response, we assumed the nonlinear subunit takes the form of a DivS model,

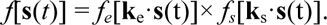

Similar to parameter estimation of the DivS models of synaptic current response, we alternately estimated the filters and nonlinearities until they converged. The same set of constraints was applied to the excitatory and suppressive nonlinearities.

### Quantification of contrast adaptation with LN analysis

We performed a more traditional LN model analysis to gauge the adaptation to contrast of both the observed data as well as the predictions of nonlinear models, following (Chander and Chichilnisky, 2001). We first separately performed LN analysis on each contrast level. The resulting nonlinearities were then aligned by introducing a scaling factor for the x-axis and an offset for the y-axis. The associated scaling factor was incorporated into the linear filters such that contrast adaptation effects are attributable entirely to changes in the linear filter.

Once the linear filters at both contrasts were obtained, we calculated contrast gain as the ratio of standard deviations of the filters at low and high contrast conditions. To make more detailed comparisons about the filter shape, we also calculated a biphasic index, based on the ratio of the most negative to the most positive amplitude of the LN filter k, i.e., |min(k)/max(k)|.

### Evaluation of model performance

We fit all models on the 7-sec segments of unique stimuli out of each 10-sec block, and cross-validated model performance on the 3-seconds repeat trials. We calculated the predictive power, or percent of explainable variance (Sahani and Linden, 2003), to quantify how well the model captured the trial-averaged response for both intracellular and extracellular recordings. This metric corrects for noise-related bias due to a limited number of trials. Note that for validation of spike-based models, we simulated individual instances of spike trains using a non-homogeneous Poisson process, and the model predictions were based on many repeats for which we generated a PSTH. All measures of model performance compared predicted to measured responses using 1-ms bins, which was necessary to measure how accurately the different models captured temporal precision (Butts et al., 2011; Butts et al., 2007).

### Coherence analysis of synaptic current response

The general model performance metrics such as predictive power and cross-validated likelihood do not reflect which aspects of the response are not captured by the model. We thus devised a new coherence-based metric to quantify how well the model performs across frequencies. The coherence between the model predicted current response *c(t*) and the recorded current response on the *i*th trial 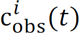 is (Butts et al., 2007):

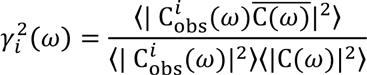

where C(ω) and 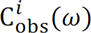 are the Fourier transforms of c(*t*) and 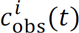 respectively, and the bar denotes complex conjugate. We used angular frequency ω = 2π*f* instead of *f* to be consistent with common conventions. The coherence measure on individual trials was averaged across repeats for each cell.

Because the observed response on each trial contains noise, a coherence of one throughout the frequency is not a realistic target. To correct for this bias, we calculated the coherence between the trial-averaged current response (i.e., the ideal predictor of response) and the recorded current on each trial. This noise corrected coherence metric represents an upper bound of coherence that can be achieved by any stimulus-processing model. It also reflects the consistency of current response at each frequency range. For example, in the low contrast condition, the response contained little high frequency components (Fig. 7A-B), and consequently the measured coherence was close to zero above 30 Hz.

### Covariance analysis of synaptic current response and spike train

We performed covariance analysis on both synaptic current responses and spike trains. Spike-triggered covariance analyses followed established methods (Fairhall et al., 2006; Liu and Gollisch, 2015; Samengo and Gollisch, 2013; Schwartz et al., 2006), Briefly, for spike-triggered covariance analysis, we collected the stimulus sequence *s_n_*(τ) = *s*(*t_n_* − *τ*) that preceded each spike time *t_n_*, where the lag τ covers 200 time bins at 1 ms resolution. The spike-triggered average was calculated as the average over all spikes 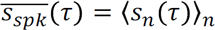, and the covariance matrix was calculated as 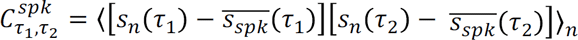. The prior covariance matrix was also computed using the above formula, but for all times rather than spike times, and it was subtracted from 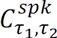 and the result diagonalized matrix to obtain its eigenvalues and corresponding eigenvectors.

This procedure was extended to perform the same analysis on synaptic currents, following past applications (Fournier et al., 2014). For current-based covariance analysis, we calculated the cross correlation between stimulus and synaptic current response (analogous to the spike-triggered average) 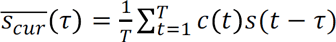, and the current-based covariance matrix was given by 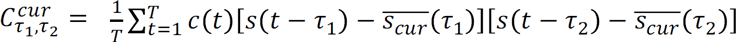. We again subtracted an average covariance matrix (unweighted by the current) and calculated eigenvalues and eigenvectors for the result.

We generated response predictions of the current-based covariance model using the two eigenvectors with largest magnitude, and applying the methods to fit a two-dimensional nonlinearity described above. The performance of the resulting model is reported in Fig. 2E, and example fits are shown in Supplemental Figure 2. To be applied to spike trains, such methods require much more data, and we could not generate firing rate predictions of the spike-based model with reasonable accuracy given the limited data to estimate a two-dimensional nonlinearity, consistent with previous applications of spike-triggered covariance to retina data (e.g., (Liu and Gollisch, 2015)). Note that simply estimating separate one-dimensional nonlinearities for each filter (e.g., (Sincich et al., 2009)), results in significantly worse predictive performance (e.g., (Butts et al., 2011)), due to the non-separability of the resulting nonlinear structure, as well as the inability for such analyses to factor in spike refractoriness.

### Event analysis of spike trains

We modified a previously established method to identify short firing episodes (events) in the spike train (Berry et al., 1997; Butts et al., 2010; Kumbhani et al., 2007). Specifically, events were first defined in the peristimulus time histogram (PSTH) as times of firing interspersed with periods of silence lasting ≥ 8 ms. Each resulting event was further analyzed by fitting the PSTH with a two-component Gaussian mixture models. An event was broken into two events if the differences of means of the two Gaussian components exceed two times the sum of standard deviations. Event boundaries were defined as the midpoint between neighboring event centers and were used when assigning event labels to simulated spikes. Events were excluded from further analysis if no spike was observed on more than 50% of the trials during the event window. This criterion excluded spontaneous spikes that occur on only a few trials. Event analysis was first performed on responses at high contrast. Events at low contrast were defined using the event boundaries obtained from high contrast data. These particular methods were chosen because they gave the most reasonable results with regards to visual inspection, but the results presented here do not qualitatively depend on the precise methods.

Once events were parsed, we measured several properties associated with each event relating to their precision and reliability (Figs. 1 and 6). First, we measured the jitter in the timing of the first-spike, using the SD of the first spike of the event on each trial. The event time scale is estimated as the SD of all spike times in each event, which is related to the duration of each event. The event Fano factor measures the ratio between the variance of spike count and the mean spike count in each event.

### Statistical tests

All statistical tests performed in the manuscript were non-parametric Wilcoxon signed rank tests, unless otherwise stated. All significant comparisons were also significant using t-tests.

## ACKNOWLEDGEMENTS

This work was supported by NSF IIS-1350990 (YC, DAB) and NIH EY021372 (YVW, SJHW, JBD).

---

**Supplemental Figure 1.**
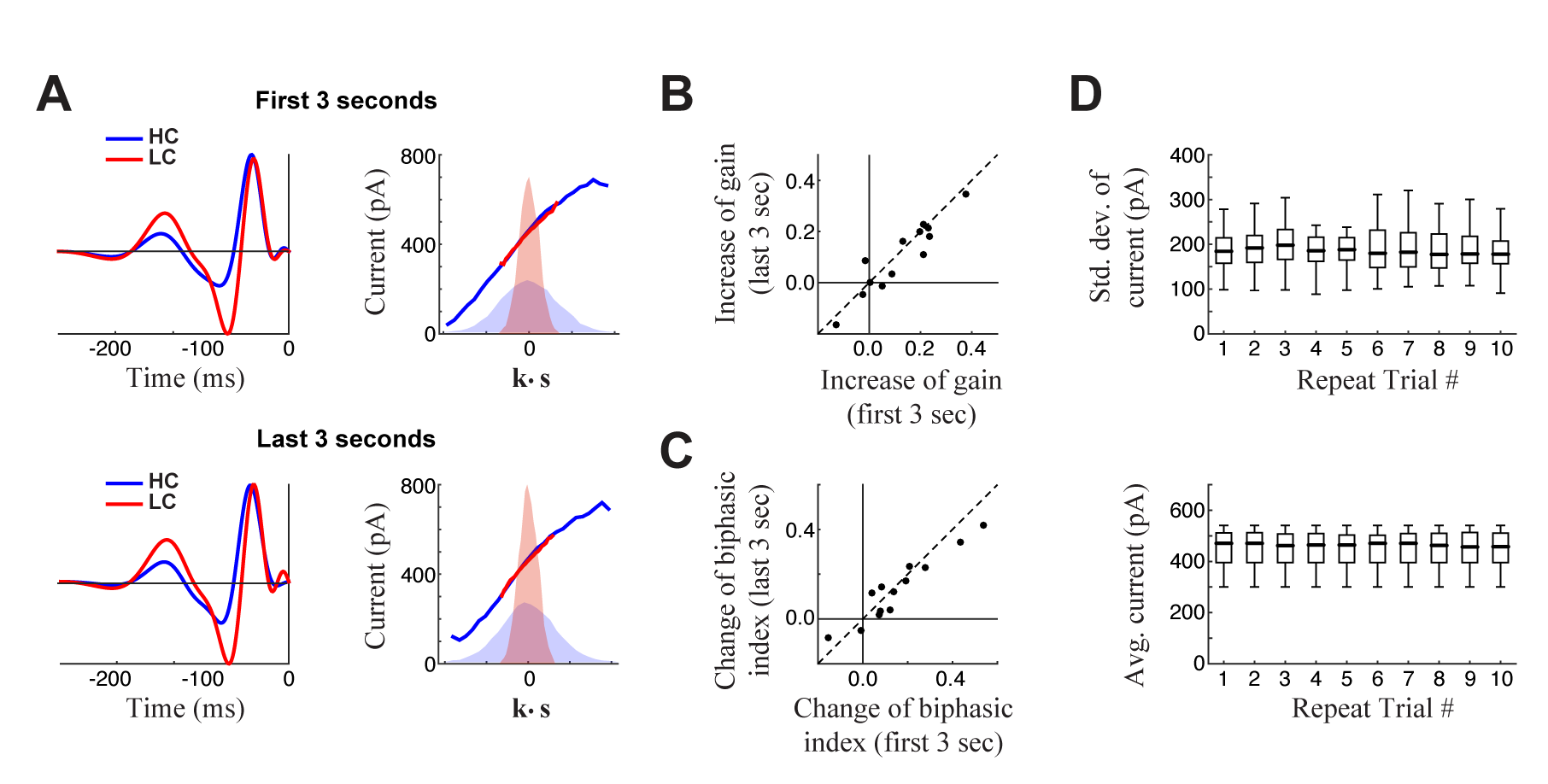
Absence of slow contrast adaptation and stability of recording. **(A-C)** We tested the presence of slow contrast adaptation by performing separate LN contrast adaptation analysis (e.g., Chander and Chichilnisky, 2001) based on data at the beginning and end of each 7-sec period of unique responses within a 10-sec trial (excluding the final 3-sec period of a repeated stimulus). **A.** LN analyses for the first 3-seconds (*top*) and last 3-seconds (*bottom*) of the trial. Model filters (*left*) are computed and scaled such that the nonlinearity (*right*) is shared between high contrast (HC, blue) and low contrast (LC, red) models, shown for an example cell. The models are nearly identical, suggesting there is no slow contrast adaptation over the course of the trial. (**B-C**) Change of contrast gain (B) and biphasic index (C) across contrasts measured on the first 3 seconds (*x-axis*) and on the last 3 seconds (*y-axis*) of each trial across neurons (*n*=13). We observed no difference on a cell-by-cell basis. **D**. To test stability of the recording, we calculate the standard deviation of intracellular synaptic current responses (*top*) and average current (*bottom*) over the 10 trials of each experiment (box plots show data aggregated across cells, *n*=13). This demonstrates that recordings are stable across trials.

**Supplemental Figure 2.**
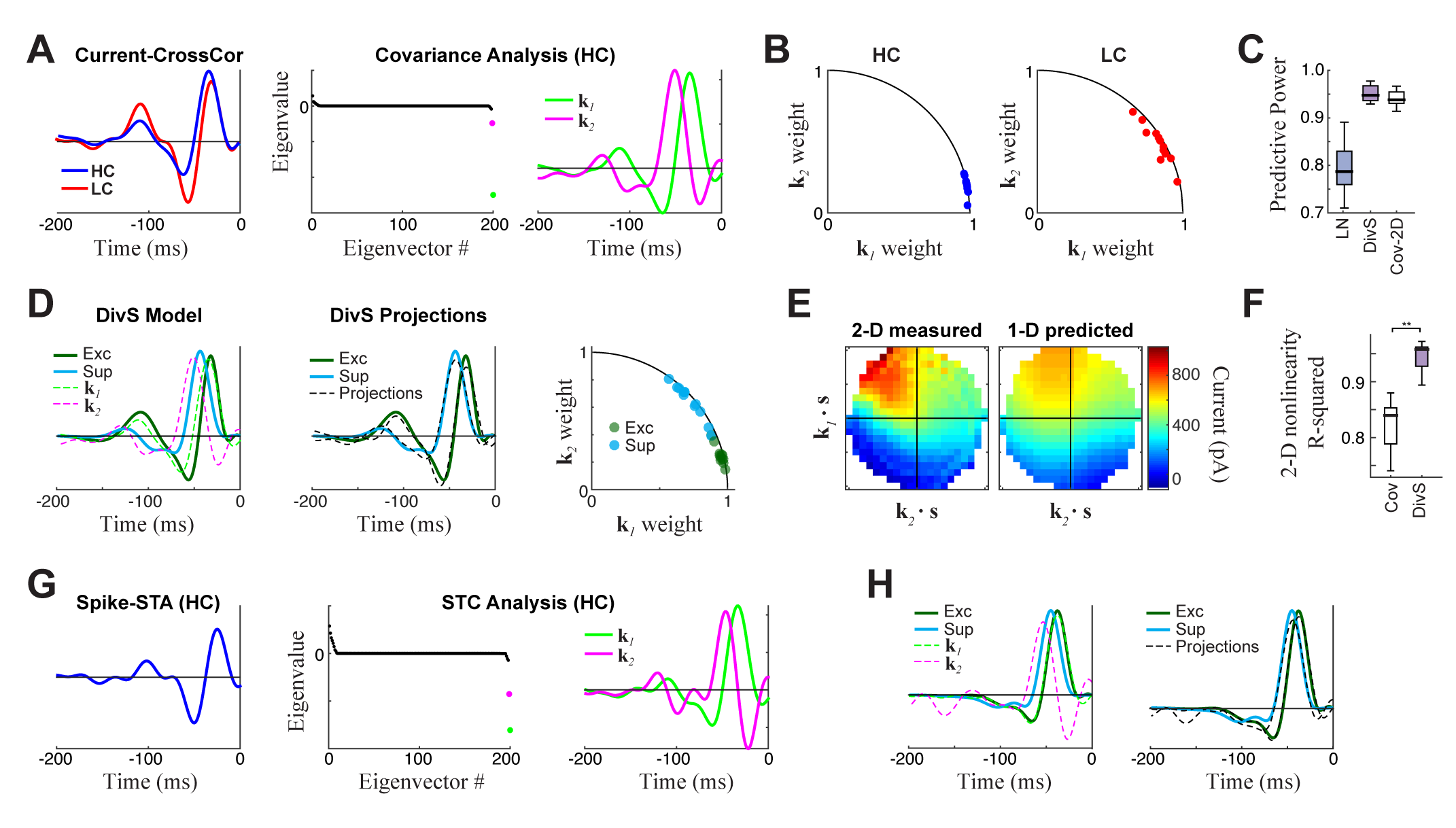
Comparison to covariance-based models. (A-F) Covariance-based analysis of synaptic currents for the same example cell as in Figure 1. Covariance analysis follows the intuition of spike-triggered covariance, but uses continuous current input rather than spikes (see Methods). **A.** *Left:* Cross-correlation between the stimulus and current response (the equivalent of a spike-triggered average) for high contrast (HC, blue) and low contrast (LC, red) stimuli. Filters are scaled to have the same standard deviation, for comparisons of shape. *Middle:* The eigenvalue spectrum for the response-triggered covariance matrix in HC, revealing two significant eigenvalues (color-coded). *Right:* The corresponding eigenvectors. **B**. The locations of the cross-correlations in HC (blue, *left)* and LC (red, *right)* within the 2-D subspace spanned by the two significant eigenvectors for all neurons («=13). Because they are all close to the unit circle, both HC and LC cross-correlations are largely contained in the covariance (COV) subspace, consistent with previously reported results for spikes (Liu and Gollisch, 2015). **C.** Model performance for the LN, DivS, and COV models («=13), reproduced from Figure 2E. This demonstrates that the COV filters coupled to a 2-D nonlinearity (described below) can nearly match the performance of the DivS model. **D.** *Left:* The excitatory (green) and suppressive (cyan) filters of the DivS model, plotted in comparison to the filters identified by covariance analysis (dashed lines). *Middle:* The DivS model filters share the same 2-D subspace as the covariance filters, as shown by comparing the filters to optimal linear combinations of the COV filters (black dashed), following previous work based on spikes (Butts et al., 2011). *Right:* The DivS filters project into the COV filters subspace across neurons, using the same analysis as in (B). Their proximity to the unit circle shows they are almost completely in the covariance subspace for all neurons, again consistent with previous work with spikes (Butts et al., 2011). **E.** *Left:* The 2-D nonlinearity associated with the COV filters, for the example neuron considered. *Right:* The best 2-D nonlinearity reconstructed from 1-D nonlinearities operating on the COV filters. Unlike the 2-D nonlinearity associated with the DivS filters (Fig. 2F), this nonlinearity cannot be represented as the product of two 1-D nonlinearities. **F.** The separability of 2-D nonlinearities for the COV and DivS models, measured as the ability of the 1-D nonlinearities to reproduce the measured 2-D nonlinearity *(R^2^)* across neurons (**p<0.0005, «=13). **(G-H)** An example neuron for which there was enough spiking data to perform a meaningful spike-triggered covariance (STC) analysis (see Methods). **G.** The spike-triggered average (*left*), eigenvalue spectrum *(middle)*, and significant STC filters *(right)*. **H.** As with the analyses of current responses above, the DivS filters (green, cyan) did not match those identified by STC (*left*, dashed), but were largely contained in the subspace spanned by the STC filters (*right*), as shown by comparing to their projections into the STC subspace (dashed black). Note that there was not enough data to estimate 2-D nonlinearities for the spiking data, and so no comparison of STC model performances could be made.

**Supplemental Figure 3.**
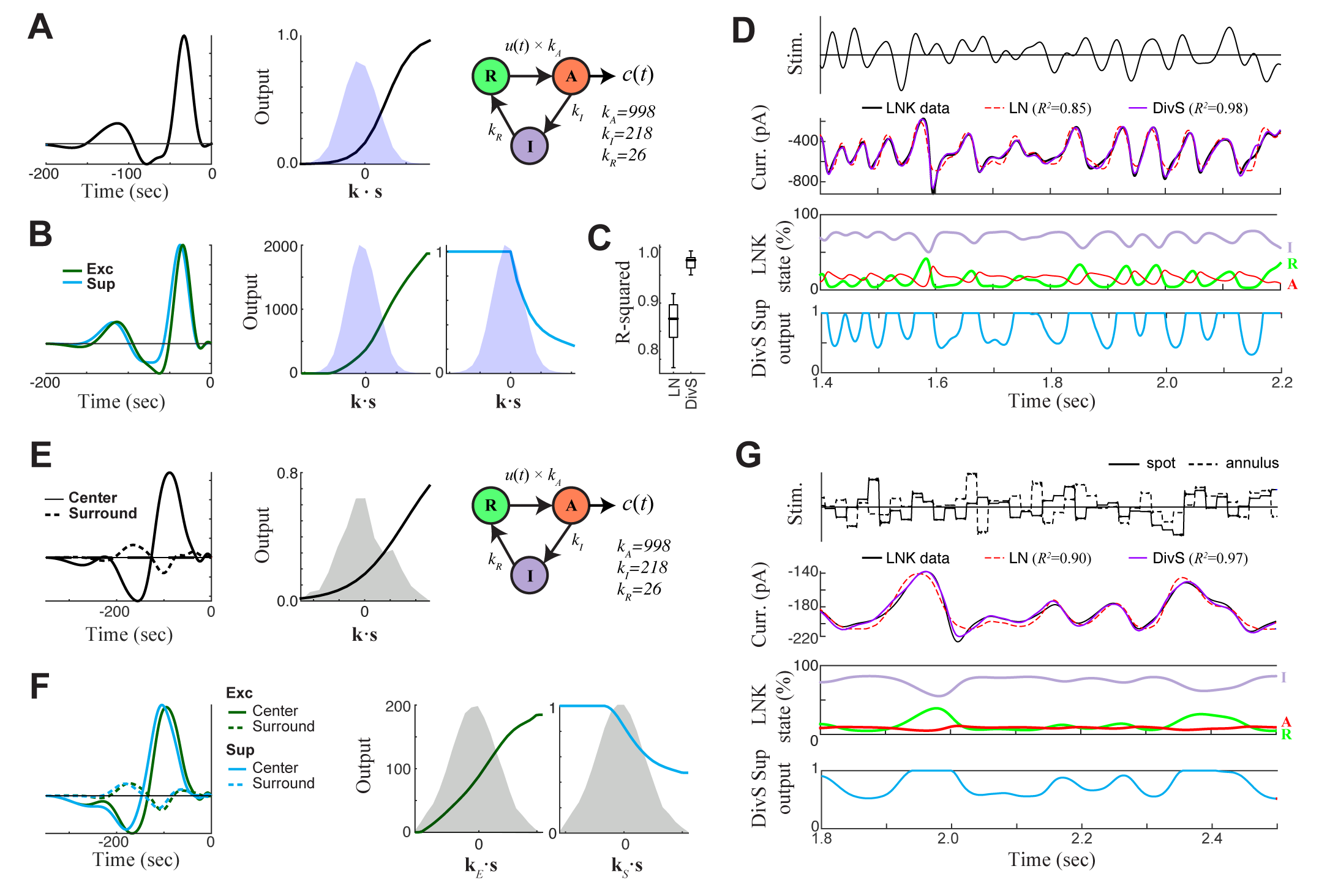
DivS model localizes the suppressive components of LNK model and reproduces its simulated response. We simulated LNK models resembling the example neurons considered in Figure 4. (**A-D**) LNK simulation in response to temporally modulated spot. **A**. The LNK model components consist of a temporal filter **k** *(left)* and static nonlinearity *f*(•) *(middle)*, whose output *u*(*t*)= *f*[**k** · **s**(*t*)] governs the transition rate between the resting (R) and active (A) states. Current output is proportional to active state occupation, and other constants govern transition to inactive (I) state and back to resting state. Parameters for this LNK simulation were derived from an LNK fit to an example neuron (see Methods). **B.** A DivS model was fit to the LNK model simulated response, with components labeled as in Figure 2. The temporal filter of suppression (cyan) is delayed relative to excitation *(left)* and only results in suppression for ON stimuli, as expected given its relationship to synaptic depression. **C**. Model performance *(R*^2^) for the LN model and DivS model across all neurons demonstrates that the DivS model can reproduce LNK simulations with greater than 90% accuracy, across simulations of all LNK models of recorded neurons (*n*=13). **D.** Simulated response of the LNK model in (A) in response to a temporal modulated spot stimulus *(top). 2nd row:* The output of the LNK simulation (black) can be reproduced better by a DivS model (red) fit to the simulated data, as compared to the LN model (blue). *3rd row:* The occupation of each internal state determines the current output in addition to the output of the LN component of the model. *4th row:* The dynamics of the divisive suppression of the DivS model (cyan) roughly match the occupation of the resting state of the LNK model (*3rd row*, green): the resting state occupancy (and availability for transition to the active state and resulting current output in the LNK model) becomes low at the same times there is suppression in the DivS model. (**E-G**) LNK simulation in response to the spot-annulus stimulus. **E.** LNK components are labeled identically as in (A), but now the filter **k** consists of separate components for the spot (*left*, solid) and annulus (dashed) regions of the stimulus. The temporal filter and nonlinearity were derived from the example cell in Figure 4B, but the kinetics parameters of the temporally modulated stimulus (A) were used in place of those derived for the spot-annulus stimuli, since the latter parameters did not result in nonlinear effects. **F**. A DivS model was fit to the LNK model simulated response, and components labeled as in Figure 2, resulting in the expected delayed ON suppression (as with the temporally modulated spot simulations in B). **G**. Simulated response using the LNK model with the spot-annulus stimulus, again with the divisive suppression of the DivS model (*4th row*, cyan) capturing the occupancy of resting state of the LNK model (*3rd row*, green).

**Supplemental Figure 4.**
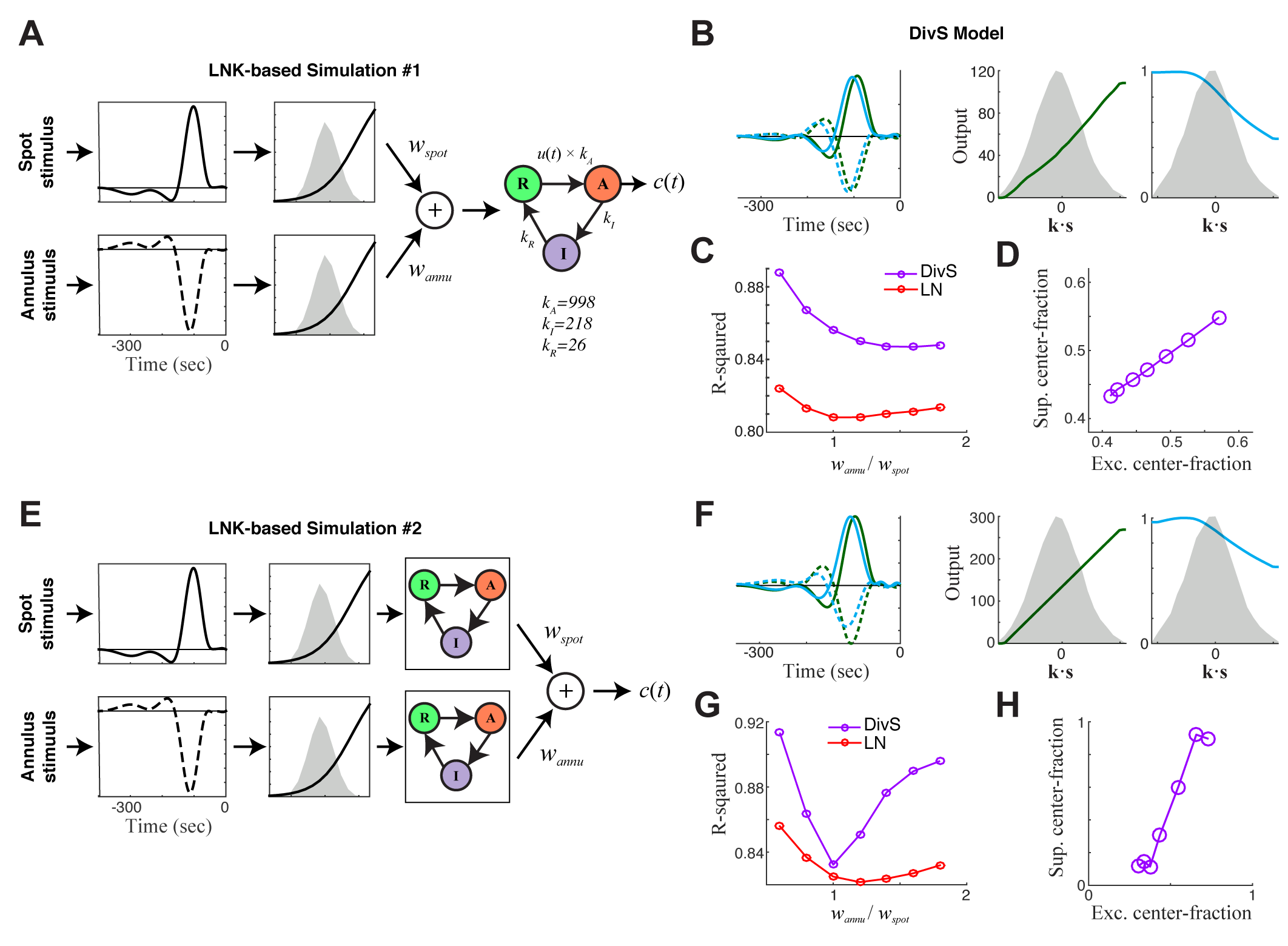
DivS model descriptions of extended LNK models. Here we consider additional model structures involving synaptic depression. The simulations here incorporate nonlinear rectified subunits, and are limited to two components corresponding to those independently modulated in the stimulus: spot and annulus. (**A-D**) First we considered an extended LNK model with independent stimulus processing of the spot and annulus stimuli, and a shared synaptic depression stage. **A.** Model schematic, showing that the separate “center” and “surround” components (corresponding to spot and annulus stimuli) are each rectified before being combined, and fed into the LNK model for synaptic depression, using the same kinetic parameters considered for simulations in Supplemental Figure 3. Simulated data was generated for a range of models of this form, where the weight for the ‘spot’ component *W_spot_* was fixed and the annulus component weight *w_annu_* was varied. **B**. The DivS model components fit to an example simulated response of the extended LNK model (with *w_spot_* = *W*_annu_). As with simpler circuits (e.g., Suppl. Fig. 3), suppression is delayed relative to excitation. Note that the DivS model was limited to only a single rectified component to match the form used to describe experiments described in Figure 4. **C**. The performance of the LN (red) and DivS (purple) models across simulations with different annulus component weights. The DivS model is significantly better than the LN model over a wide range of parameters (each point corresponds to the results of simulation with different choice of *w_annu_*), suggesting a large portion of the synaptic depression effect is captured by the DivS model. Note, however, that the DivS model has a harder time explaining this [simulated] data than the data from real ON Alpha ganglion cells (i.e., Fig. 4D). **D.** For all simulations, the “spatial profile” of suppression matched that of excitation, as measured by the “center fraction”, which was given by the norm of the center component of the filter divided by the norm of the full filter. [The center fraction is 1 if for no surround component, and zero for no center component.] (**E-H**) We next considered an extended LNK model with both independent stimulus processing and independent kinetics. **E.** Model schematic, showing the separate center and surround components each with independent synaptic depression — again with the same kinetic parameters previously considered. **F**. The DivS model components fit to an example simulated response of the extended LNK model (with *W_spot_* = W_annu_). **G**. Performance of DivS model and LN model on simulated LNK model response. **H**. Tight correlation of the center fractions of excitation versus divisive suppression of the DivS model components (as in panel D). Over this and other types of simulations involving synaptic depression, we never observed the case where DivS excitation is largely from the center and suppression is largely from the surround (e.g., Fig. 4E).

